# Pervasive morphological responses to climate change in bird body and appendage size

**DOI:** 10.1101/2023.09.28.560061

**Authors:** Sara Ryding, Alexandra McQueen, Marcel Klaassen, Glenn J. Tattersall, Matthew R.E. Symonds

## Abstract

Changes to body size and shape have been identified as potential adaptive responses to climate change, but the pervasiveness of these responses is questioned. To address this, we measured body and appendage size from 5013 museum bird skins of ecologically and evolutionary diverse species. We found that morphological change is a shared response to climate change across birds. Birds increased bill surface area, tarsus length, and relative wing length through time, consistent with expectations of increasing appendage size in warmer climates. Furthermore, birds decreased in absolute wing length, consistent with the expectation of decreasing body size in warmer climates. Interestingly, these trends were generally consistent across different diets, foraging habitats, and migratory and thermoregulatory behaviours. Shorter-term responses to hot weather were contrary to long-term effects for appendages. Overall, our findings support that morphological adaptation is a widespread response to climate change in birds that is independent of other ecological traits.

## Introduction

Among endotherms, there are ecogeographical clines in body and appendage size, called Bergmann’s^1^ and Allen’s rule^2^, respectively. These rules state that animals in warmer climates tend to be smaller in size, with relatively larger appendages. The rules are said to facilitate improved thermoregulation by increasing the body’s surface area to volume ratio^3^. Support for Bergmann’s and Allen’s rules has been found in many endotherms^4, 5, 6, 7, 8^, with thermal adaptation appearing to best explain the trends^4, 9^. The provenance of ecogeographical gradients in morphology, and the physiological mechanisms that could underpin these, has led to the hypothesis that animals will respond to anthropogenic climate change with morphological adaptation, as a temporal extension of the ecogeographical rules^10^.

Appendage size is predicted to follow an Allen’s rule-style response to climate change, becoming larger relative to body size as climate warms. Increases in relative appendage size have been called ‘shape-shifting’^11^, due to their effect on overall body shape. Examples of shape-shifting responses to climate change have been identified in some birds^12, 13, 14, 15^ and mammals^16, 17, 18^, including changes in bills, legs, and ears. However, most studies have focussed on a limited range of species, meaning we don’t know how pervasive these changes are, or whether morphological changes vary according to species’ characteristics.

Understanding the impact of species characteristics is key for bird bills, which have well-established links to diet and habitat^19^ that may affect their shape-shifting. The consistency of Allen’s rule across different foraging types^4^ may lead us to predict that morphological adaptation to climate change will not differ between diet categories, but even small changes in bill morphology can influence diet^20^ which may constrain adaptation to climate change. Other ecological traits (e.g. movement and behaviour) may also affect the ability to adapt. For example, Allen’s and Bergmann’s rule are more pronounced in resident compared to migratory species^4, 7^, which may influence shape-shifting responses. Similarly, there are relationships between appendage size and use of thermoregulatory behaviours across^21^ and within species^22^, meaning such behaviours may also affect adaptive responses over time. While Allen’s rule is explained by thermal adaptation to different climates^4, 7^, the multifaceted role of appendages means we have been unsure how ecological traits affect shape-shifting across diverse species.

Bergmann’s-style body size changes in mass are a commonly identified morphological response to climate change, having been primarily documented in birds^23, 24, 25, 26, 27, 28, 29, 30, 31, 32^. Body mass decreases are consistent across birds with different diets^25, 26^, migratory behaviours^24^, and degrees of human commensalism^26^, thus supporting body size decreases as a generalizable response to climate change in birds^33^. While long-term responses of body mass to climate change seem consistent, some studies find that body mass can increase in the short-term (i.e. contrary to long-term trends) in response to warm weather^25^. Contrary long-term and short-term trends may imply that immediate responses to short-term weather conditions reflect foraging^25^ and reproductive capacity^32^, rather than overarching adaptation to climatic warming. Long-term data spanning more than a few decades could help disentangle short-term weather phenomena from the effect of cumulative climate change.

One challenge with ascertaining changes in body size is that there are several alternative ways of measuring it. In birds, absolute wing length is highly correlated with body mass^34^ and underlying bone length^35^ and has therefore been considered an indicator of structural body size that is more stable than mass^33, 36^. Indeed, many studies on Bergmann’s rule find that absolute wing length decreases with lower latitudes^8^. However, the response of absolute wing length to climate change is not consistent (e.g.^26^). Different multi-species comparisons have identified decreases^24, 37^, increases^23, 25, 36^, or no changes^26, 28, 32^ in absolute wing length through time. Increases in absolute wing length over time is contrary to how body size has been predicted to respond to climate change^10^, but have been attributed to mechanisms like enabling more efficient flight^23, 25^ or a response to subtle changes in habitat structure^25^. Testing wing length responses in a broader clade of species, who use different types of habitats, can help us understand what causes disparate responses in wing length to both long-term climate change and short-term weather.

Another proposed reason that wing length may increase in response to climate change is the potential role of the wing as a heat-shedding appendage^31^. When considering Allen’s rule in birds, it is often the bills and tarsi that are measured and considered appendages^5, 38^. This is because these appendages are unfeathered, highly vascularized, and have a demonstrated role in thermoregulation, making them obvious locations for heat loss^39, 40^. The undersides of wings are also highly vascularised and have been proposed to form another appendage which may conform to Allen’s rule^31, 41^, although the role of wing vascularity in shedding heat is poorly known (but see^42, 43, 44^). However, if the wing functions as a heat-shedding appendage, we expect wing length relative to body size to increase in an Allen’s rule-style response (*sensu* ^31^), rather than absolute wing length. It is possible, therefore, that studies finding decreases in absolute wing length have missed increases in relative wing length by not considering body mass as a covariate.

We investigated morphological responses of Australian birds to climate change. Using a dataset of over 5000 individual museum specimens from the past 100 years, representing 78 species from 14 diverse orders, we examined changes in appendage and body size in response to climate change and short-term weather conditions. We measured bill surface area and tarsus length as metrics of appendage size, and absolute wing length as a body size indicator. Using a subset of 2000 birds from 73 species, we further investigated changes in relative wing size, as an indicator of appendage size, and body mass, as an indicator of body size. We examined whether morphological adaptation to climate change following Allen’s and Bergmann’s rule was a shared response across diverse avian lineages. To investigate whether constraints imposed by diet or nutritional stress could be mediating morphological responses, we tested whether responses differed between diet categories. Similarly, to test whether morphological responses are mediated by behaviour, we assessed whether migratory and thermoregulatory behaviours affected changes through time. Lastly, to better understand how responses to short-term weather may differ from long-term trends and interact with habitat, we investigated if use of different foraging habitats influenced morphological change to short-term weather.

## Methods

We measured 5013 museum bird skins collected over the past century. These represented 78 different species from 14 orders with contrasting diets, foraging habitats, thermal behaviours, and movement ecology (Table S1), ranging from 31 – 73 individuals per species (average 64). Species were chosen based on availability in the collections of the Melbourne Museum, the Australian National Wildlife Collection, the South Australian Museum, and the Australian Museum. We focused on species with good coverage through the past century (median sampling period beginning 1906 and ending 2010), where “good coverage” was defined as species with at least 10 individuals collected every 2 decades throughout the time period. We chose 2014 as the maximum cut-off year to eliminate the possible effect of post-collection museum shrinkage, which only occurs in the first years following skinning^45^. We selected individual specimens based on collection year (to ensure a good spread through time for each species) and age at collection (only adults, as determined by tag information on age or skull ossification, were measured). We chose specimens within species from a similar location, to minimize the influence of geographic clines in morphology (i.e. Bergmann’s and Allen’s rule on spatial scales) on our detection of temporal changes in morphology.

### Morphological measurements

We used absolute wing length and body mass as indicators of body size to investigate decreasing body size through time. To investigate appendage size increases, we measured bill surface area, tarsus length, and wing length relative to body mass. For all 5013 individuals, we measured bill surface area and wing length. Bill surface area was measured using 3D scans taken with a structured light 3D surface scanner (Artec Space Spider, Artec 3D, Luxemburg). In Artec Studio 14, meshes were registered and outliers pixels (points whose mean distances were more than 3 standard deviations away from other points) were removed, before 3D models were generated using the ‘sharp fusion’ tool. The bill feather line was traced using the ‘eraser’ tool to define the bill boundaries, and the bill surface area was subsequently measured using the ‘sections’ function. Wing length was measured as the length of the flattened wing cord, using a butted ruler. For 5010 individuals, we also measured tarsus length using digital callipers (Whitworth). We obtained bird mass at time of collection from museum tags when available (n=2213). Collection data for all 5013 birds was obtained from the museum data repository Ozcam (Online Zoological Collections of Australian Museums; https://ozcam.org.au/) and supplemented with information on the collection tags, which included time of collection (year, and when available, month and day), location (latitude and longitude), and, when available, sex.

### Responses to climate change and weather

Australia has warmed by 1.4°C on average since 1910^46^, including maximum temperature increases of 0.06°C per decade and minimum temperature increases of 0.12°C per decade^47^. We used year as a proxy for cumulative climate change over the past century, allowing for an overarching evolutionary trajectory throughout our 106-year sampling period to be detected. We were also interested in how shorter-term weather fluctuations impacted morphology. For each specimen latitude and longitude, we calculated weather conditions, averaging them over the 5 years preceding collection from a 5k gridded dataset of meteorological observations of Australia^48^, dating back to 1910 for temperature and 1900 for precipitation. Weather conditions included minimum winter and summer temperature, maximum winter and summer temperature, and summer and winter precipitation. For specimens collected prior to 1910, we used the first available 10-year average for that site to back-calculate estimated temperature conditions, since there is high autocorrelation in weather conditions from year to year for an estimated 25 years^49^. We condensed these weather conditions into a single covariate using principal component analysis, with PC1 explaining 74% of the variation (Figure S1). High values of weather PC1 were associated with higher maximum and minimum winter temperatures and higher maximum summer temperatures in the five years prior to collection.

To examine whether aspects of ecology could underlie, facilitate, or offset morphological adaptation, we collected information on diet, migratory behaviour, thermoregulatory behaviour, and foraging habitat (table S1). Diet was categorised as per EltonTraits^50^ and defined as omnivorous, fruit/nectar, invertebrate, plant/seed, and vertebrate/fish/scavenger feeders, according to the food source that constituted the majority of their diet. If there was not one single major diet source, defined as when several diet sources each contributed less than 50% of the total diet, the species was designated as an omnivore. Migratory behaviour was split into three categories; fully migratory, partially migratory, or non-migratory. Information on migration in passerines was taken from McQueen et al.^51^, and we followed the same method (i.e. assessing range maps in the Handbook of Birds of the World^52^) to score non-passerines. Any uncertainties were verified by consulting the Handbook of New Zealand and Australian Birds (HANZAB^53^). Presence (1) or absence (0) of thermoregulatory behaviours (bill tucking; tucking the bill under back feathers, unipedal roosting; standing on one leg while the other is tucked into belly or breast plumage, and sitting; covering both legs in plumage by sitting) was taken from Pavlovic, Weston, and Symonds^21^ and was available for 69 species (Table S1). Lastly, predominant foraging habitat was determined as aerial-, ground-, vegetation-, and water-based foraging, according to the percentage time spent in each foraging habitat as per EltonTraits^50^. Vegetation-based foraging was an amalgamation of understory, mid-high, and canopy foragers, whereas water-based foraging combined below water, on water surface, and pelagic foragers. For some species, the foraging habitats were evenly shared across two categories, or the foraging habitat EltonTraits listed represented the predominant habitat where prey was caught rather than where birds spent most of their time foraging. For these species, we consulted HANZAB^53^ and determined the species categorisation based on the detailed descriptions (see Table S2).

### Statistical analysis

All statistical analysis was carried out in R version 4.1.1^54^.

To investigate whether morphology was changing through time and with weather, we ran linear mixed models under a Bayesian framework using the ‘brms’ package^55^. To account for the phylogenetic relatedness of species in our dataset, we constructed a phylogeny by downloading 1000 trees from the BirdTree website (www.birdtree.org^56^) using the Hackett backbone^57^ and the ‘phangorn’ package^58^ to generate the maximum clade credibility tree with the maxCladeCred function. The phylogenetic matrix was included as a random effect in all following models. To account for measures of multiple individuals per species, we included species identity as a random effect. All analyses were run for 100 000 iterations, with a 10 000 burn-in period and thinning set to 1000.

Morphological measures were scaled and centred prior to analysis (i.e. *z*-transformed), which made the measures comparable across diverse species. All models can be found in table S3. Each morphological measure (bill surface area, tarsus length, wing length, and body mass) was the response variable in models where year and PC1 of the weather data (“weather PC1” from here on) were the fixed effects (table S3), each with weak uninformative priors (mean of 0, with variance of 5 standard deviations). Year was centred on 1964, but not scaled, prior to analysis. While weather generally became warmer through time, the correlation between weather PC1 and year was not strong enough (r=0.03) to prevent including both predictors in the same model. This also indicates that shorter-term weather fluctuates greatly compared to longer term climate trends. For models where bill surface area and tarsus length were the response variable, we included wing length as a fixed effect to account for body size variation (table S3). Because appendage size is likely to increase with increasing body size, we gave wing length a weakly informative prior in models where bill and tarsus size were the response variable (mean of 1, with variance of 5 standard deviations). Absolute wing length is considered a good indicator of body size^34^, hence it is a suitable metric to correct for size and evaluate changes in relative bill and tarsus size, and results in a larger sample size than using body mass as the metric for body size. To investigate whether *relative* wing length changed through time and in relation to weather, we included body mass as a fixed effect for one model where wing length was the response variable (table S3), with a weakly informative prior (mean of 1, with variance of 5 standard deviations). Lastly, to see whether increases in appendage size were due to concomitant decreases in body size, we tested for changes in absolute bill and tarsus size by running these models without controlling for body size (i.e., without wing length; table S3).

To examine whether species had different responses to climate change, we ran models of bill, tarsus, and wing size including species as a fixed effect (rather than random effect), with an interaction with year (table S3). To encourage the model to converge, the priors for this were stronger but still uninformative (mean of 0, with variance of 2 standard deviations). This model also included the phylogenetic matrix. The sample sizes within species were small (average n = 64), meaning we might not expect significant interactions, but this would allow us to identify species which were having particularly strong responses to climate change. We could not do this for relative wing size or mass, because our sample size within species was too small (mean n = 30).

We further examined ecological traits that might influence the presence and strength of any morphological changes. To test whether adaptation differed according to ecological traits, we expanded the previous models to include diet category, migratory behaviour, thermoregulatory behaviours, and foraging habitat (table S3), each with weak uninformative priors (mean of 0, with variance of 5 standard deviations). Diet category and foraging habitat were strongly correlated and therefore excluded from the same models. To investigate whether ecological variables affected morphological adaptation through time, we included an interaction between year and diet category, year and migratory behaviour, and year and each of the thermoregulatory behaviours (table S3).

Lastly, we investigated whether foraging habitat influenced morphological responses to short-term weather. To this end, we included an interaction between weather PC1 and foraging habitat, but still including year as a separate fixed effect in the model (table S3). We examined the interaction estimates to ascertain whether different categories were significantly different from each other, and then extracted the marginal effects for each category using the ‘emmeans’ package^59^.

## Results

### Responses to cumulative climate change

We found widespread support for morphological adaptation to climate change in several morphological measures. Across species, there were consistent increases in relative bill surface area, relative tarsus length, and relative wing length through time (Figure 1). The increase in bill size, tarsus length and relative wing length were 0.0002, 0.0004, and 0.0003 standard deviations per year, respectively (table Extended Data 1). On average, appendages increased by 1.5% (range: –20 –19.5%), 1.8% (range: –14 – 12%) and 1.6% (range: –13.5 – 32%) for bill surface area, tarsus length, and relative wing length, respectively, over the study period. Morphological changes varied between species, where 63% and 69% of species had positive estimates of bill and tarsus size through time, respectively (Figure 2). Of these, 9 and 14 species had credible intervals entirely above 0 for bill and tarsus size change, respectively. The increases in bill and tarsus size across species were also consistent when considered in absolute terms and when including mass (rather than wing length) as our body size indicator (table ED2).

**Figure 1.**
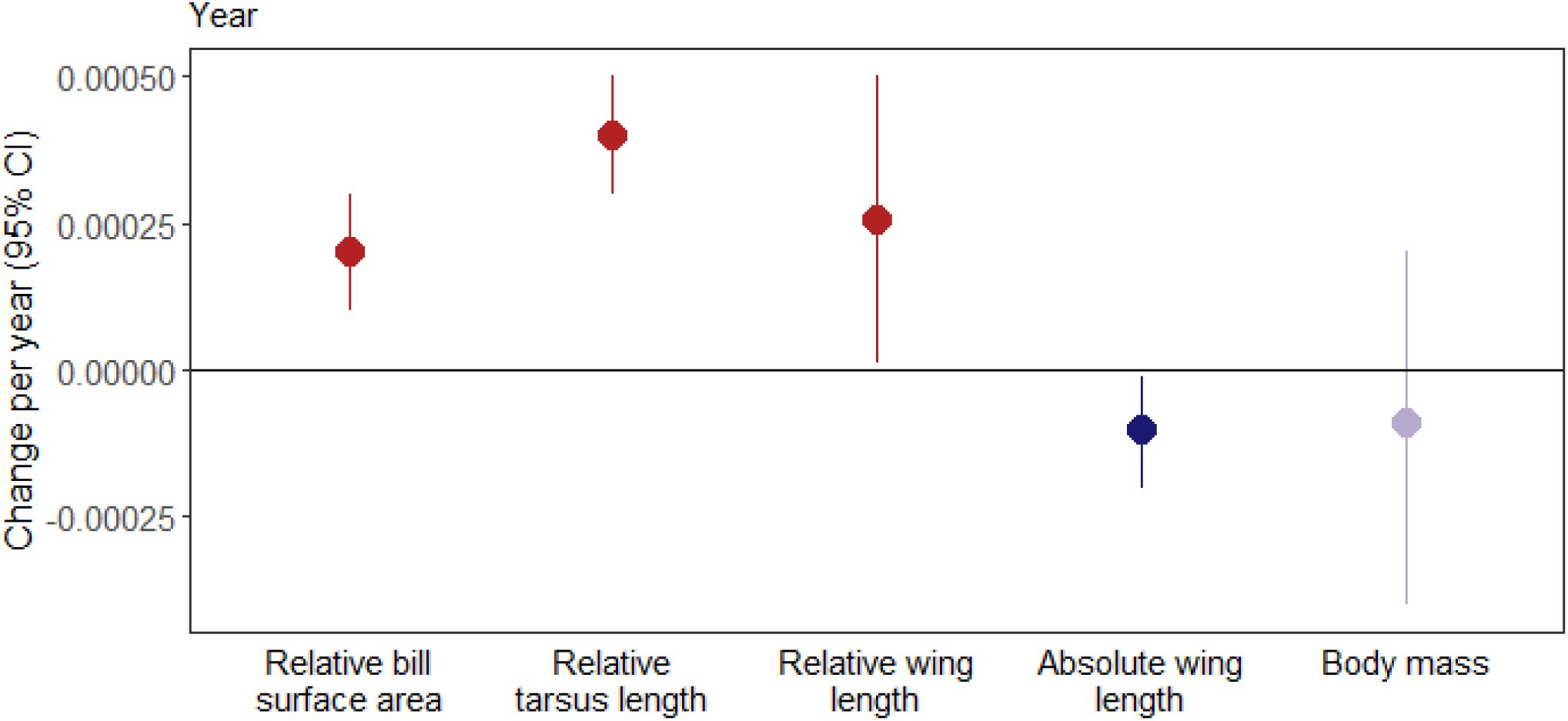
Change per year in relative bill surface area, relative tarsus length, relative wing length, absolute wing length, and body mass in Australian birds. Plot shows effect estimate and 95% credible intervals of change per year, expressed as standard deviations. Color and shade indicate mean estimates and credibility interval (CI) range, where red indicates positive mean estimates and dark red (above 0) or light red (crossing 0) indicates CI ranges. Blue indicates negative mean estimates and dark blue (above 0) or light blue (crossing 0) indicates CI ranges.

**Figure 2.**
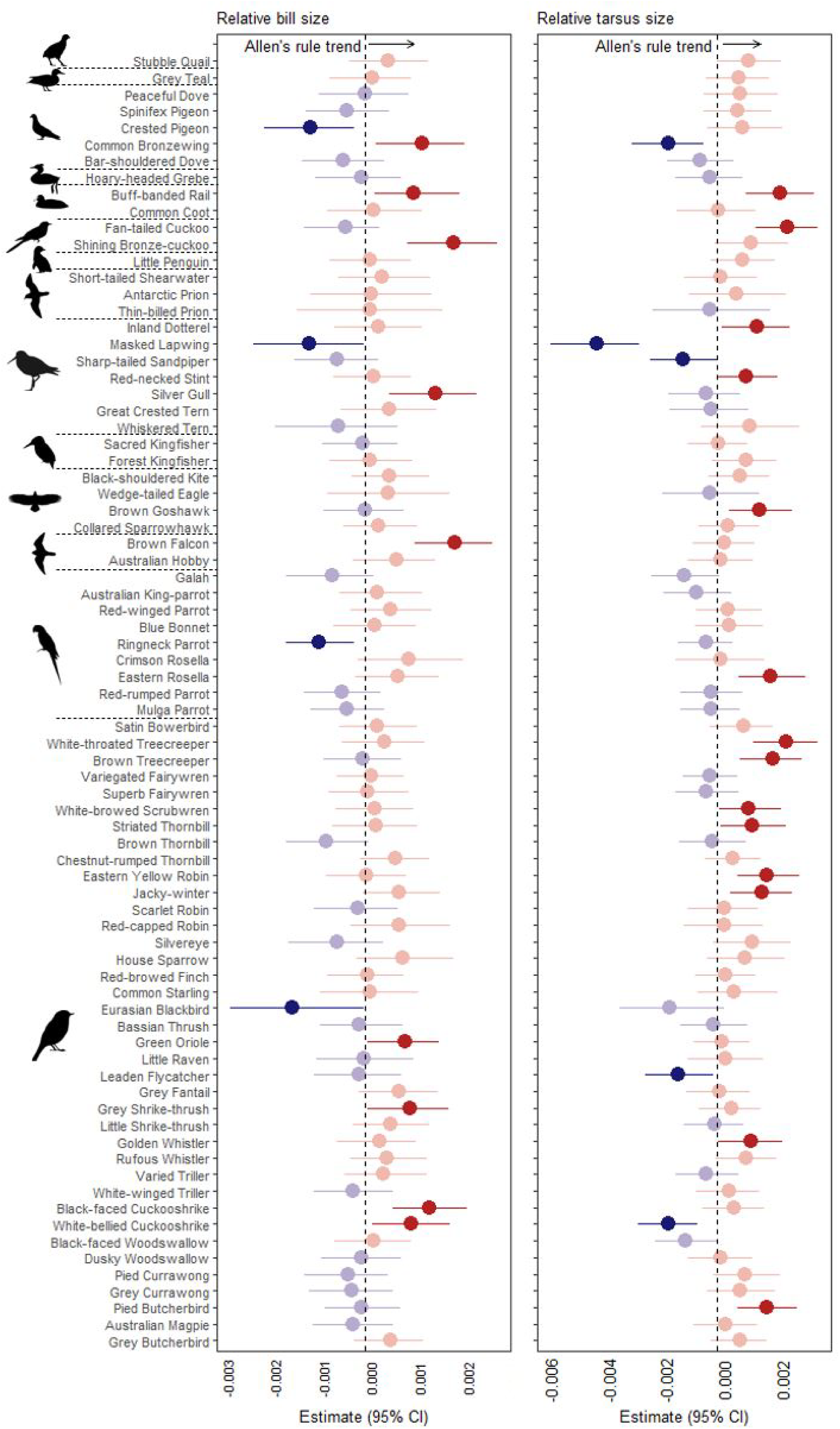
Species responses to climate change in terms of bill and tarsus size. Color and shade indicate mean estimates and credibility interval (CI) range, where red indicates positive mean estimates and dark red (above 0) or light red (crossing 0) indicates CI ranges. Blue indicates negative mean estimates and dark blue (above 0) or light blue (crossing 0) indicates CI ranges. The top arrow indicates the overall direction of change, as per Figure 1 and as expected by Allen’s rule. Species are ordered taxonomically, going by order: Galliformes, Anseriformes, Columbiformes, Podicipediformes, Gruiformes, Cuculiformes, Sphenisciformes, Procellariiformes, Charadriiformes, Coraciiformes, Accipitriformes, Falconiformes, Psittaciformes, and Passeriformes.

The increases in bill and tarsus size through time were largely not influenced by ecological traits. Increases in bill surface area were consistent across most diet categories. Vertebrate/fish/scavenger eaters increased bill size significantly more than plant/seed eaters (Figure ED1). The exception to increasing bill size were our two species of fruit/nectar feeders which decreased bill size (Figure ED1, Table ED1, N = 134). Invertebrate eaters increased tarsus size significantly more than plant and seed eaters, but both categories showed increases (Figure ED1, Table ED1). Apart from this, there were no significant differences in tarsus length changes between diet categories. Increases in relative wing length were also consistent across diet categories (Figure ED1). Migratory behaviour, and the presence of unipedal roosting and sitting behaviours had no effect on bill or tarsus size changes (Figures ED 3, 5, and 6). While the presence of bill-tucking behaviour had no influence on changes in bill size or relative wing length, species without bill-tucking behaviour showed stronger increases in tarsus length (Figure ED4). Species with unipedal roosting behaviour showed significantly greater increases in relative wing length (Figure ED4).

In addition to increasing appendage size, we found decreases in body size in response to climate change. Absolute wing size, a measure of body size, decreased through time, whereas mass showed no significant change through time (Figure 1). Absolute wing length decreased by 0.0001 standard deviations per year, with average decreases of 0.4% (range: –7 – 10%) over the study period. Eight species (10%) had confidence intervals which were entirely below 0, but there was considerable variation in absolute wing length changes among species (Figure 3). Going contrary to the overall effect of year on absolute wing length, 41% species had a positive estimate and most of these were from passerines.

**Figure 3.**
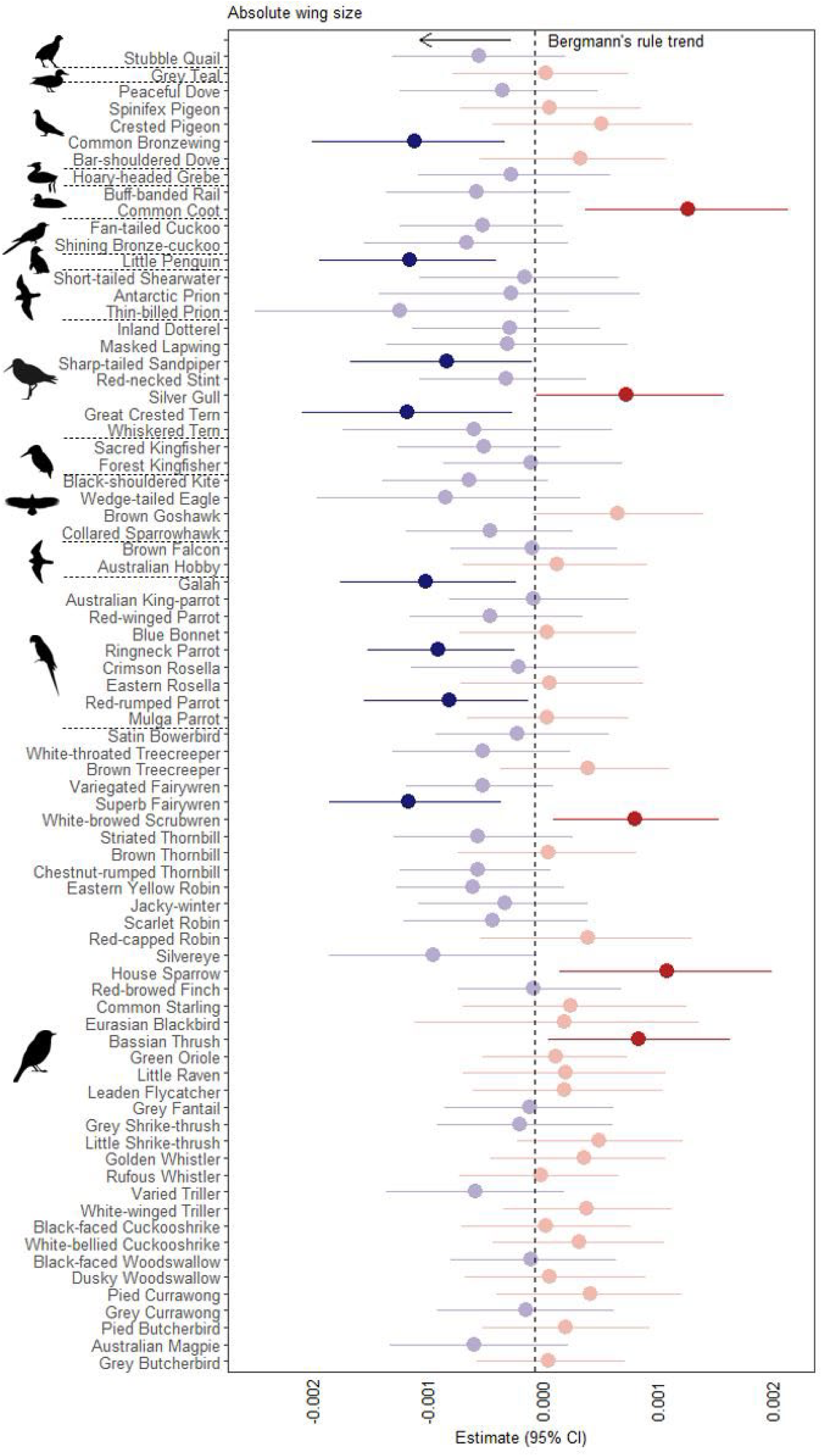
Species responses of body size (absolute wing length) to climate change. Color and shade indicate mean estimates and credibility interval (CI) range, where red indicates positive mean estimates and dark red (above 0) or light red (crossing 0) indicates CI ranges. Blue indicates negative mean estimates and dark blue (above 0) or light blue (crossing 0) indicates CI ranges. The top arrow indicates the overall direction of change, as per Figure 1 and as expected by Bergmann’s rule. Species are ordered taxonomically, going by order: Galliformes, Anseriformes, Columbiformes, Podicipediformes, Gruiformes, Cuculiformes, Sphenisciformes, Procellariiformes, Charadriiformes, Coraciiformes, Accipitriformes, Falconiformes, Psittaciformes, and Passeriformes.

The overall trend of decreasing absolute wing length varied by diet category. Plant/seed eaters and omnivores showed increases in absolute wing length through time and thus differed significantly from fruit/nectar and vertebrate/fish/scavenger eaters. Invertebrate eaters also differed significantly from vertebrate/fish/scavenger eaters (Figure ED1). Omnivores increased absolute wing length significantly more than invertebrate eaters (Figure ED1). Mass declines were generally consistent across diet categories, except invertebrate feeders increased in mass and thus significantly differed from vertebrate/fish/scavenger, omnivore, and plant/seed eaters (Figure ED1). Wing length decreases were not significantly different by behaviour (Figures ED3, 4, 5, and 6). For mass, full migrants significantly decreased in mass through time (Figure ED3), but no other behaviours moderated the relationship between year and mass.

### Responses to shorter-term weather

The responses to short-term weather PC1 were different to the long-term climate change responses. While controlling for year, we found consistent decreases in all morphological measures in response to high weather PC1, which predominantly reflected warmer winter temperatures and warmer minimum summer temperatures. Relative bill surface area, relative wing length, absolute wing length and body mass all significantly decreased with higher weather PC1, while relative tarsus length did not decrease significantly (Figure 4). The effect estimates of weather PC1 on relative bill size and relative wing length were 0.008 and 0.007, respectively. The effect estimates of weather PC1 on absolute wing length and mass were larger, at 0.01 and 0.02, respectively. Decreases in bill size and tarsus length in response to hotter weather were consistent across foraging habitats (Figure ED2). Vegetation-based foragers decreased relative wing length significantly more than water-based foragers and absolute wing length significantly more than other foraging habitats (Figure ED2). Decreases in mass were significantly larger in vegetation-based foragers compared to ground– and water-based foragers (Figure ED2).

**Figure 4.**
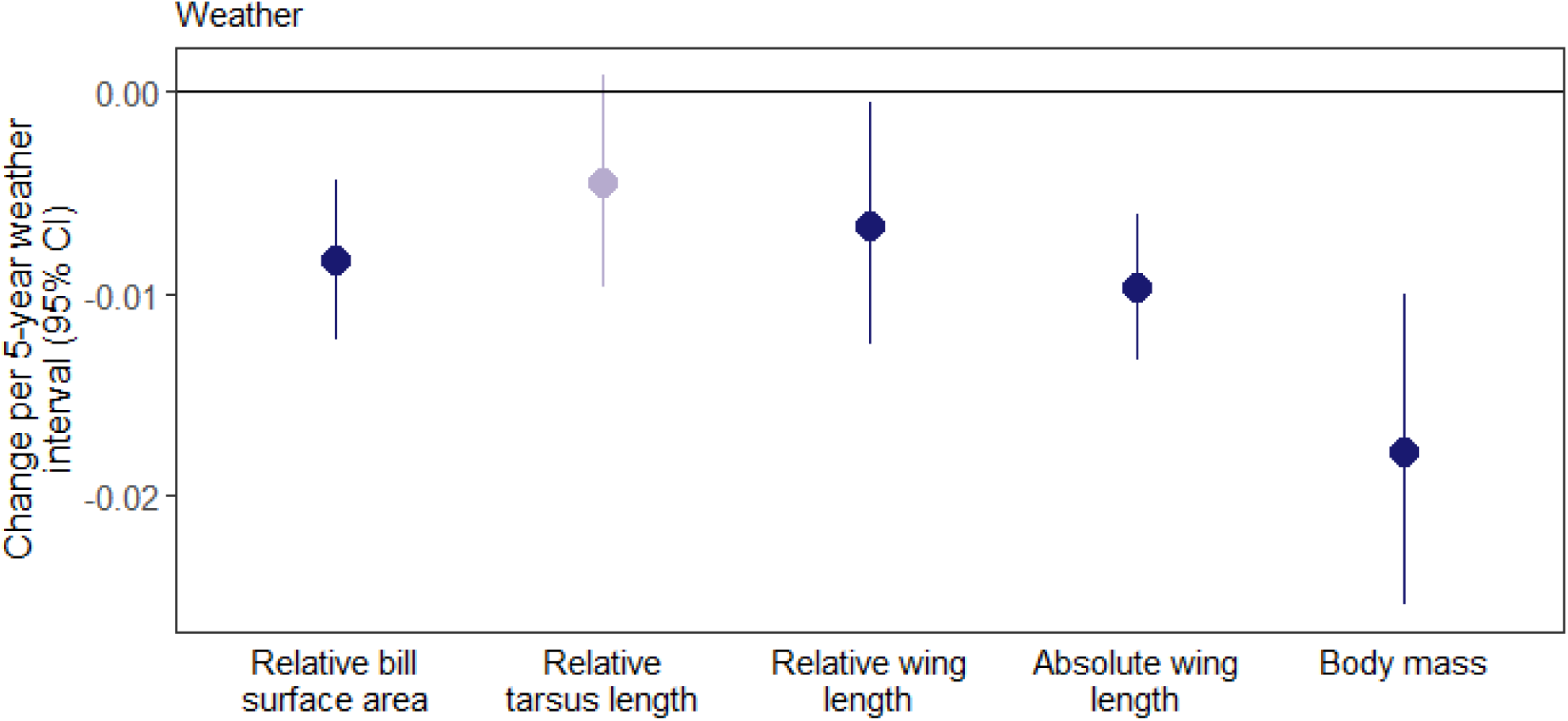
Morphological adaptation to higher weather PC1 is consistent across Australian species and occurs in almost all measured morphological metrics. Bill surface area, relative wing length, absolute wing length and mass decrease in response to hot weather. Color and shade indicate mean estimates and credibility interval (CI) range, where blue indicates negative mean estimates and dark blue (above 0) or light blue (crossing 0) indicates CI ranges.

## Discussion

We found broad evidence for increasing appendage size and decreasing body size over time, indicative of shared morphological responses to climate change in birds. These trends were largely generalisable across taxonomic orders, diet categories, foraging habitats, and behavioural suites. However, both body and appendage size decreased in response to short-term hotter weather. Collectively, these results support the premise of morphological adaptation to climate change, but indicate complex short-term selective pressures acting on body shape and size.

Relative bill and tarsus size have significantly increased over the past century. Increases in relative size of appendages (i.e. shape-shifting) are consistent with adaptive responses to climate change, wherein larger appendages facilitate improved thermoregulation^10, 11^. We found that ecological traits, such as diet, foraging habitat, and behaviour, were largely unimportant in shape-shifting, demonstrating that the response is broadly applicable across ecological traits. Bill and tarsus size increases being shared across diverse species indicates the changes are unlikely to be driven by other factors, such as nutritional stress, further supporting that morphological adaptation is a shared response to climate change in birds. Our results are consistent with other studies on appendage size change in individual endothermic species in response to climate change^12, 14, 16, 17, 18^. Overall, we estimate an increase in bill surface area of 1.5%, but up to 19% for some of our most impacted birds, which is consistent with the 4-10% increases found in bills of Australian parrots from 1871 to 2008^12^. Changes in tarsus length through time have not been explored as much as bill size (but see ^30^).

However, the adherence of tarsi to Allen’s rule^7, 38^ has led to speculation that these will respond similarly. We find tarsus length is changing more than bill surface area. Similar to the other appendages, our results show increases in relative wing length through time in response to climate change. Our results support previous research^31^, indicating an Allen’s rule-style increase in relative wing size. Overall, we show that increasing appendage size in response to climate change is shared across species and diverse ecological traits, further supporting the generality of “shape-shifting” responses.

We find decreases in absolute wing length, but not body mass, of Australian birds through time. While we do not find statistically significant decreases in body mass, the effect estimate is similar to that of absolute wing length, suggesting our lack of support for decreasing body mass may be related to the lower sample size available for mass (n = 2213, versus n = 5013 for absolute wing length). Our results can be interpreted as decreases in body size consistent with Bergmann’s rule responses to climate change^24, 33, 37^. Our results contrast with studies reporting increases in absolute wing length through time (^23, 25^, but see^31, 60^). We also find effects of diet category on changes in absolute wing length, which could indicate that nutritional stress impacting growth is driving the observed changes, and we find a phylogenetic pattern, with passerines tending to show increases in wing length compared to other groups (Figure 4). However, these effects are not consistent across other morphological metrics, and it is unclear why nutritional stress would impact passerine wing length growth, but not bill or tarsus length. Differences between diet and taxonomic order that we find cannot fully explain the differing trends found by other studies, because many of those have focused on the same diet and taxonomic group and still found contrasting results. For example, both Gardner, Heinsohn, and Joseph^37^ and Weeks et al.^23^ mostly focused on invertebrate feeding passerines, and yet found absolute wing length responses in different directions. The increase in relative, but not absolute, wing size that we find may help explain the direction in which wing length is adapting to climate change. Wing length may be under competing pressures as both an indicator of body size (via wing loading) and as a heat-shedding appendage, which may help explain the inconsistent response of absolute wing length across studies.

We find that all appendages (bill, tarsus, relative wing length) and indices of body size (mass, absolute wing length) decrease in size in response to short-term warmer weather. Declines in appendage size go against our prediction that warmer conditions would be associated with larger appendages. However, these results align with other studies which find disparate results between year and temperature^25, 27, 28^. We consider that there may be two possible mechanisms through which this discrepancy between short-term and long-term trends could occur. One mechanism for disparate short-term and long-term appendage size responses is via competing pressures imposed by thermal environment and reproduction. In the short-term, warmer weather conditions may promote greater availability of resources, thereby reducing competition and enabling less-fit individuals (i.e., smaller body sizes, and smaller appendages) to survive the winter and reproduce. This may then lead to small individuals, with smaller appendages, to become relatively more represented in the population in the short term. The short-term trend would be counter-acted by selection by the environment at life stages other than reproduction (e.g. overall survival), which would favour small-bodied but big-appendaged individuals, shifting the long-term trend back towards increasing appendage size. This has been indicated by previous research showing long-term morphological responses are not driven by reproductive fitness, but are occurring at another, unidentified life stage^32^. While some indicate that the alternative selection acting on long-term trends relates to foraging^27^, this is unlikely an important contributor in our study due to the mostly consistent results over time across diet categories and hot weather across foraging habitats. It may be that all diet categories and foraging habitats are similarly impacted by climate change, but it seems more likely that the consistent decreases in body and appendage size in response to short-term hot weather is because weather conditions affect breeding and growth in a different way compared to long-term cumulative climate change.

Another hypothesis for the contrasting trends between long-term climate and short-term weather is that, unlike in the generally cooler North America where other research was done^32, 61^, warmer weather in Australia makes development and nestling provisioning more difficult (*sensu* ^62^ in Africa). The more difficult conditions mean only the best adapted individuals (with small bodies and large appendages) survive. However, in the short-term, hot weather deleteriously impacts chick development as adults are less able to provision young, with the stunted growth resulting in smaller bodies and appendages. Like our results, hot weather 2 years before resulted in decreases in all morphological metrics in Amazonian birds, going opposite to the overall trend through time, which was ascribed to the stressful conditions caused by hot weather^25^. Furthermore, we find that vegetation-based foragers appear more impacted by hot weather than other foragers for certain morphological metrics (Figure ED2), which may indicate these species are more vulnerable to short-term hot weather and unable to provision chicks, suffering stunted growth as a result. Once conditions improve, it is these stunted individuals with genotypes for big appendages and small bodies (but current phenotypes of small bodies and small appendages) who constitute the population and reproduce. The next generation inherits big appendages and small bodies, as these are heritable traits^63, 64^. Genotypes of small bodies and large appendages could therefore become overrepresented through time, while cycles of stunted growth in response to hot weather drives disparate trends in the short term.

## Acknowledgements

We thank the Australian National Wildlife Collection (ANWC), Museums Victoria (NMV), the South Australian Museum (SAM) and the Australian Museum (AM) for access to specimen and permission for scanning. Specifically, we’d like to thank the following staff: Leo Joseph, Chris Wilson, Julian Teh, Kylea Clarke, Karen Roberts, Maya Penck, and Leah Tsang for assistance. We thank David J. Wilkinson for assistance with some bill surface area measurement extraction. We thank Kristina Macdonald, Anthony Rendall, Janet Gardner and Kalya Subasinghe for helpful discussions and statistical advice.

Funding was provided by an Australian Research Council Discovery Project (DP190101244) to M.R.E.S., M.K. and G.J.T. Research funding was also provided by a Natural Sciences and Engineering Research Council of Canada Discovery Grant to G.J.T. (RGPIN-2020-05089), and from the Centre for Integrative Ecology at Deakin University to S.R.

## Extended Data

**Figure ED1.**
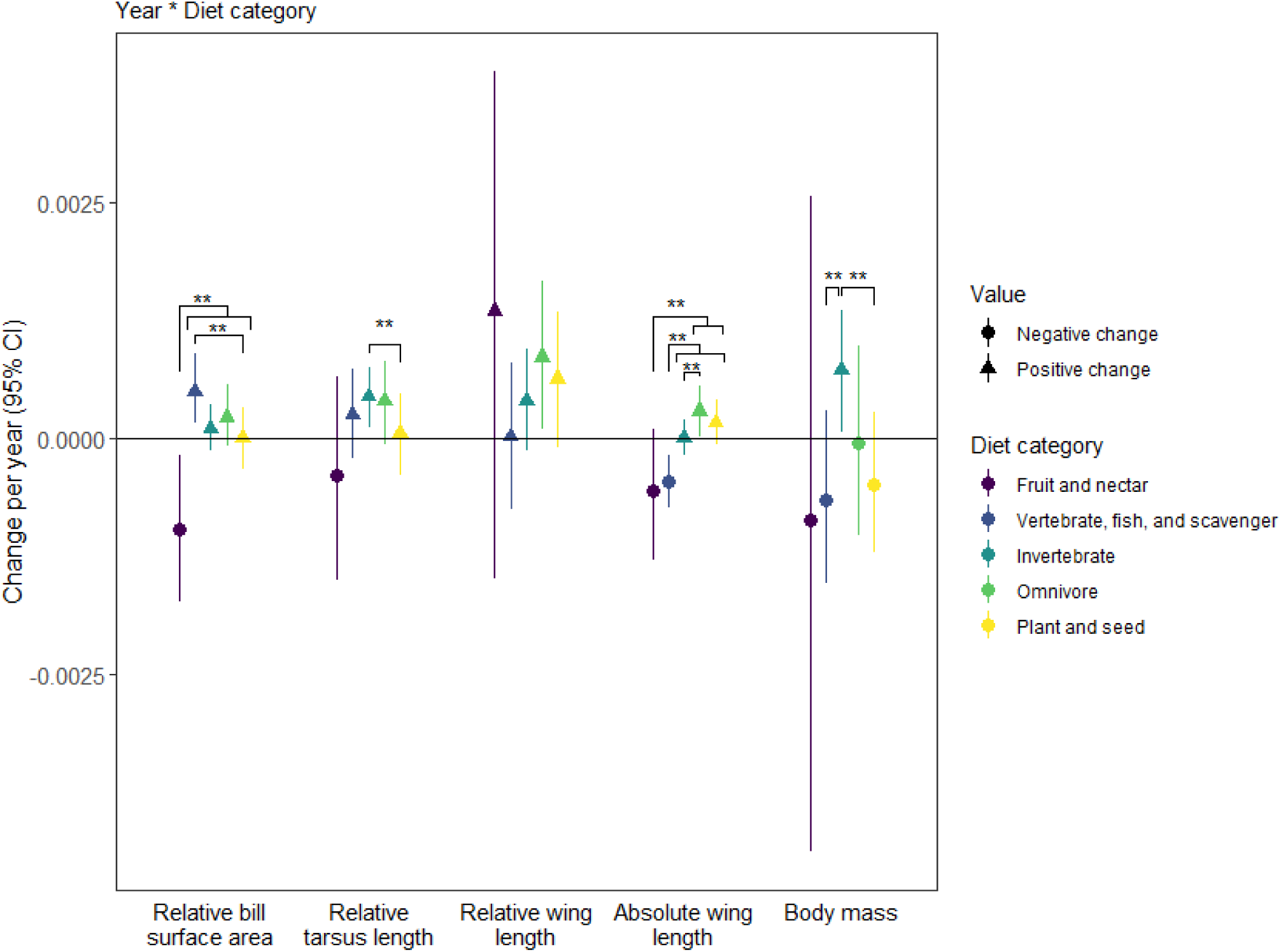
Increases in bill and tarsus size in response to year were largely consistent. The exception were fruit and nectar feeders, which decreased in size and thus significantly differ from all other diet categories for bill size. Vertebrate/fish/scavenger feeders increased bill size significantly more than plant/seed eaters, but both increased bill size through time. Tarsus size increased significantly more in invertebrate feeders compared to plant/seed eaters, but both increased tarsus size through time. Increases in relative wing length did not differ between any diet categories. Decreases in absolute wing length in response to year differed between diet categories, with fruit/nectar feeders decreasing significantly more than omnivores and plant/seed eaters and vertebrate/fish/scavenge feeders decreasing significantly more than invertebrate, omnivore, and plant/seed eaters. Omnivores increased absolute wing length significantly more than invertebrate feeders. Most groups decreased mass through time, except invertebrate feeders which differed significantly from vertebrate/fish/scavengers and plant/seed eaters. Triangles represent positive marginal effects, whereas circles represent negative marginal effects. Overall, there were increases in bill, tarsus, and relative wing size, but decreases in wing length and mass through time.

**Figure ED2.**
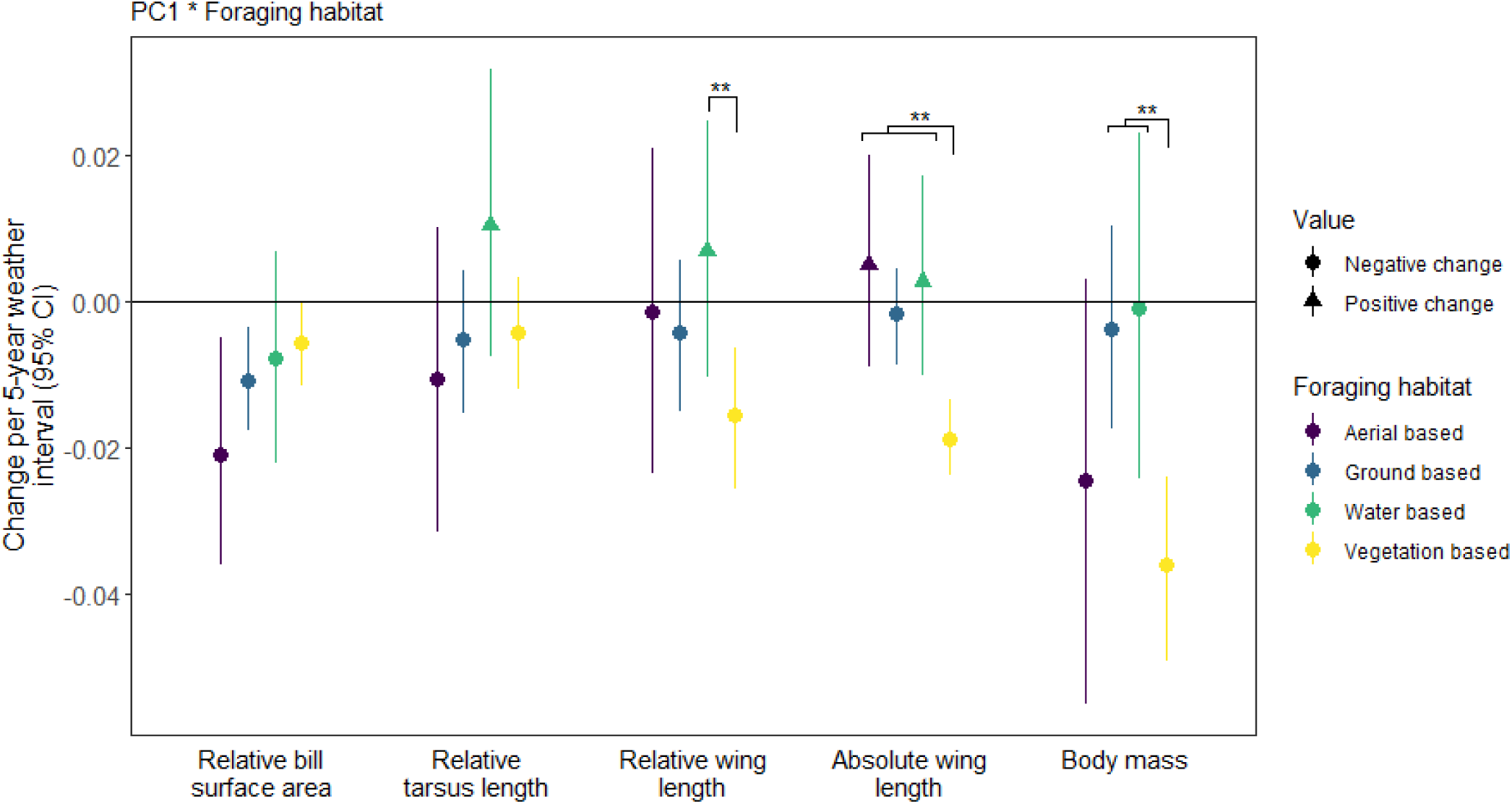
Decreases in bill surface area and tarsus length in response to weather in the 5 years preceding capture did not differ significantly between foraging habitats. Vegetation-based foragers decreased relative wing length significantly than water-based foragers. Similar patterns were seen in absolute wing length and mass; absolute wing length decreased significantly more in vegetation-based foragers compared to all other foraging habitats, whereas mass decreased significantly more in vegetation-based foragers than ground– and water-based foragers. Triangles represent positive marginal effects, whereas circles represent negative marginal effects. Overall, there were decreases in bill size, relative wing length, absolute wing length and mass in response to hotter weather.

**Figure ED3.**
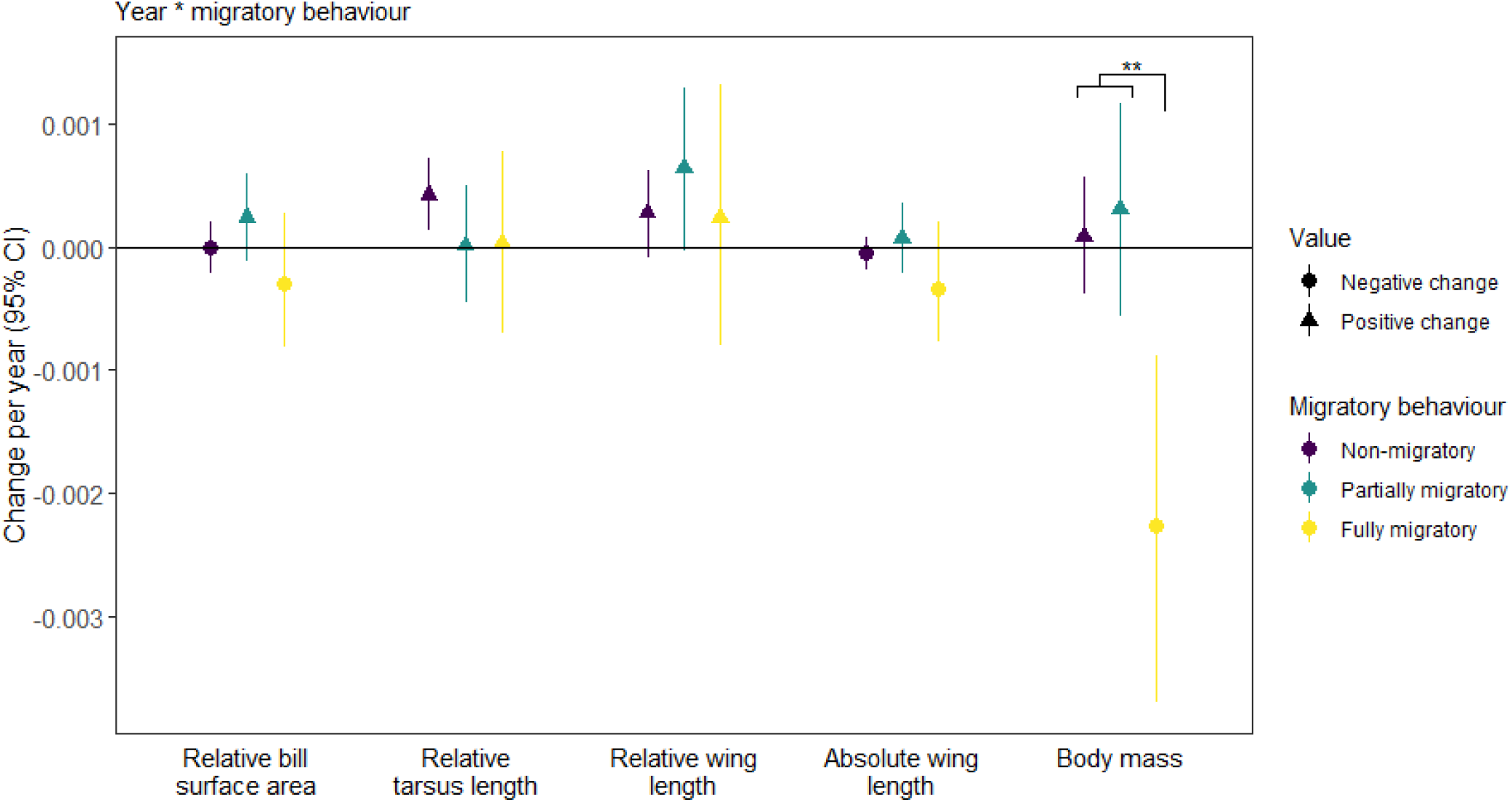
Changes in morphology through time were rarely different between migratory behaviours. The exception to this was mass, which decreased more for fully migratory birds than for non and partial migrants. Triangles represent positive marginal effects, whereas circles represent negative marginal effects. Stars signify significant differences between categories. Overall, there were increases in bill, tarsus, and relative wing size, but decreases in wing length and mass through time.

**Figure ED4.**
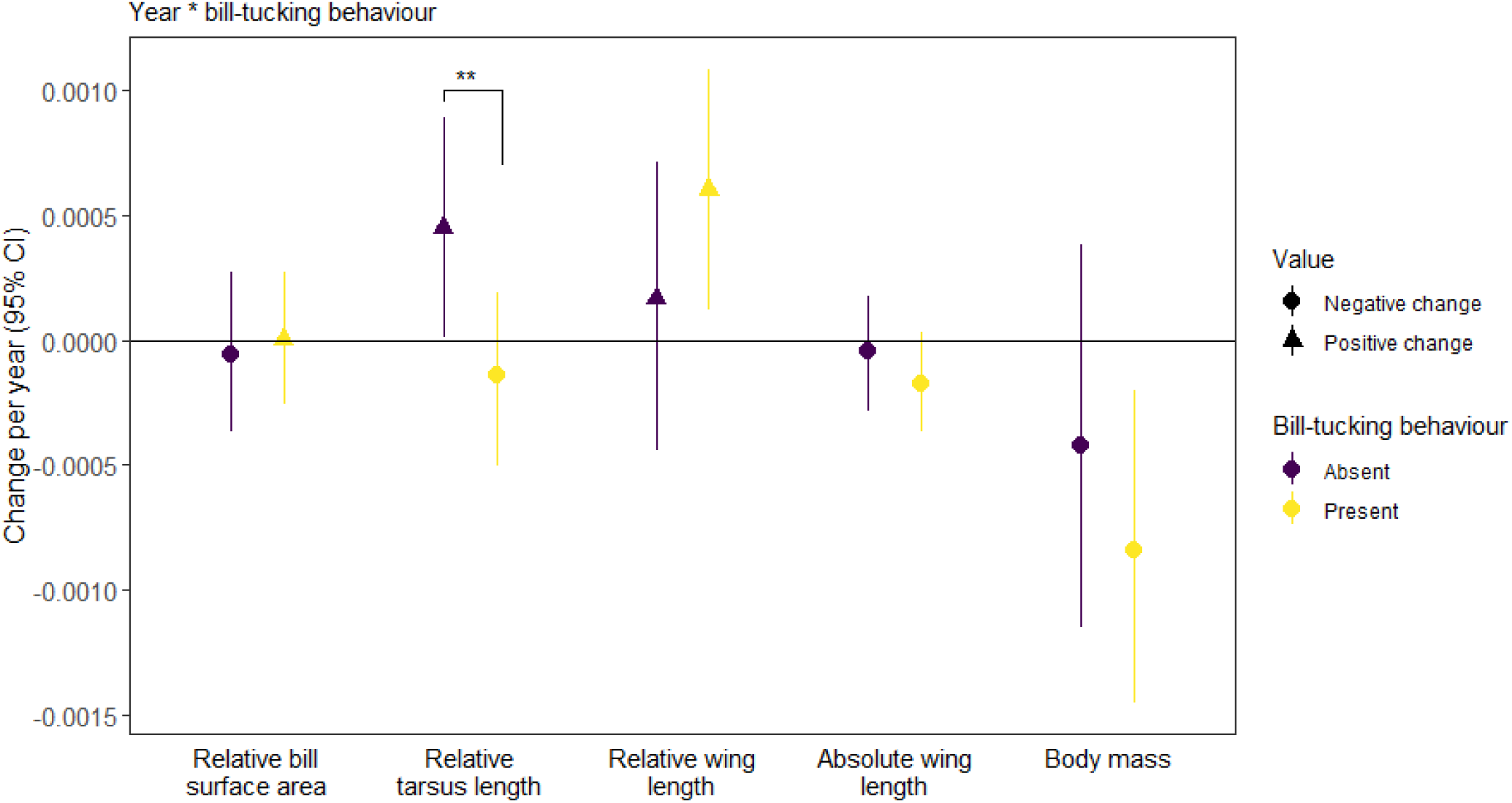
Morphological changes through time did not significantly differ between bill-tucking behaviours for bill size, relative wing length, absolute wing length, and mass. Tarsus length increased significantly more through time in birds that do not engage in bill-tucking behaviour. Triangles represent positive marginal effects, whereas circles represent negative marginal effects. Stars signify significant differences between categories. Overall, there are increases in bill, tarsus, and relative wing size, but decreases in wing length and mass through time.

**Figure ED5.**
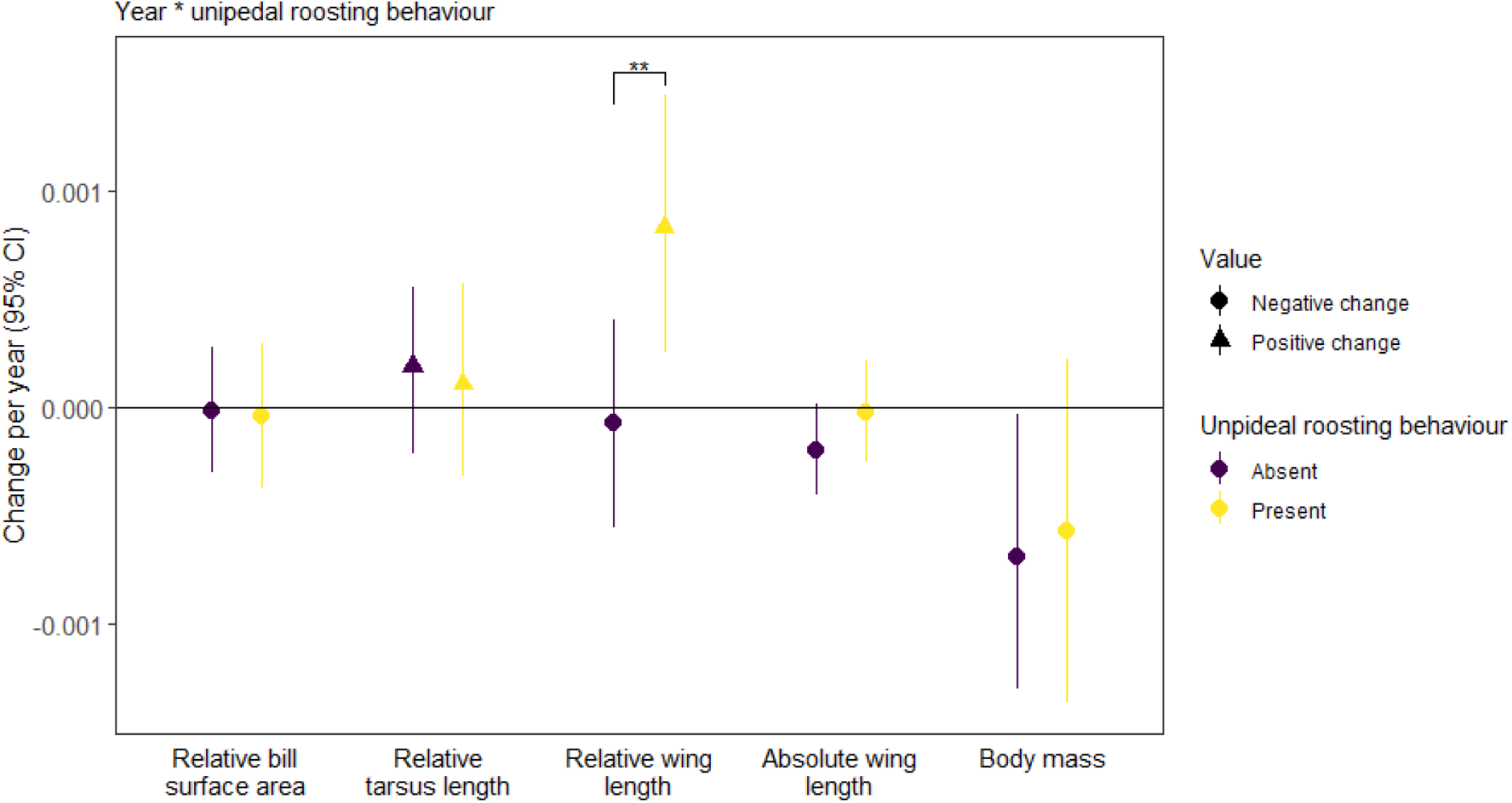
Morphological changes through time did not significantly differ between unipedal roosting behaviours for bill size, tarsus length, absolute wing length, and mass. Relative wing length increased significantly more through time in bird species that engage in unipedal roosting behaviour. Triangles represent positive marginal effects, whereas circles represent negative marginal effects. Stars signify significant differences between categories. Overall, there were increases in bill, tarsus, and relative wing size, but decreases in absolute wing length and mass through time.

**Figure ED6.**
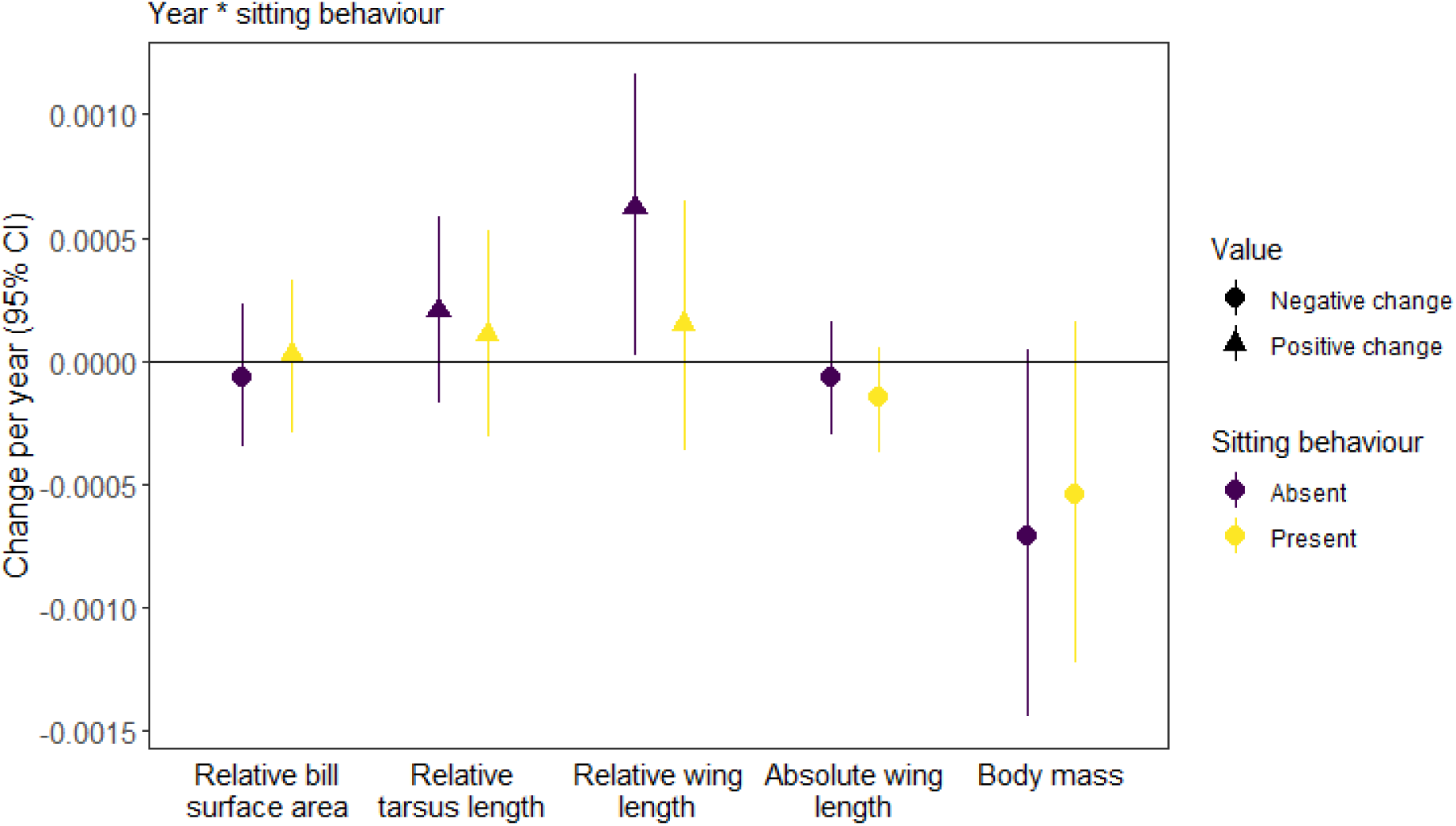
Morphological changes through time did not significantly differ between sitting behaviours for any morphological metrics. Triangles represent positive marginal effects, whereas circles represent negative marginal effects. Stars signify significant differences between categories. Overall, there were increases in bill, tarsus, and relative wing size, but decreases in wing length and mass through time.

**Table ED1.**
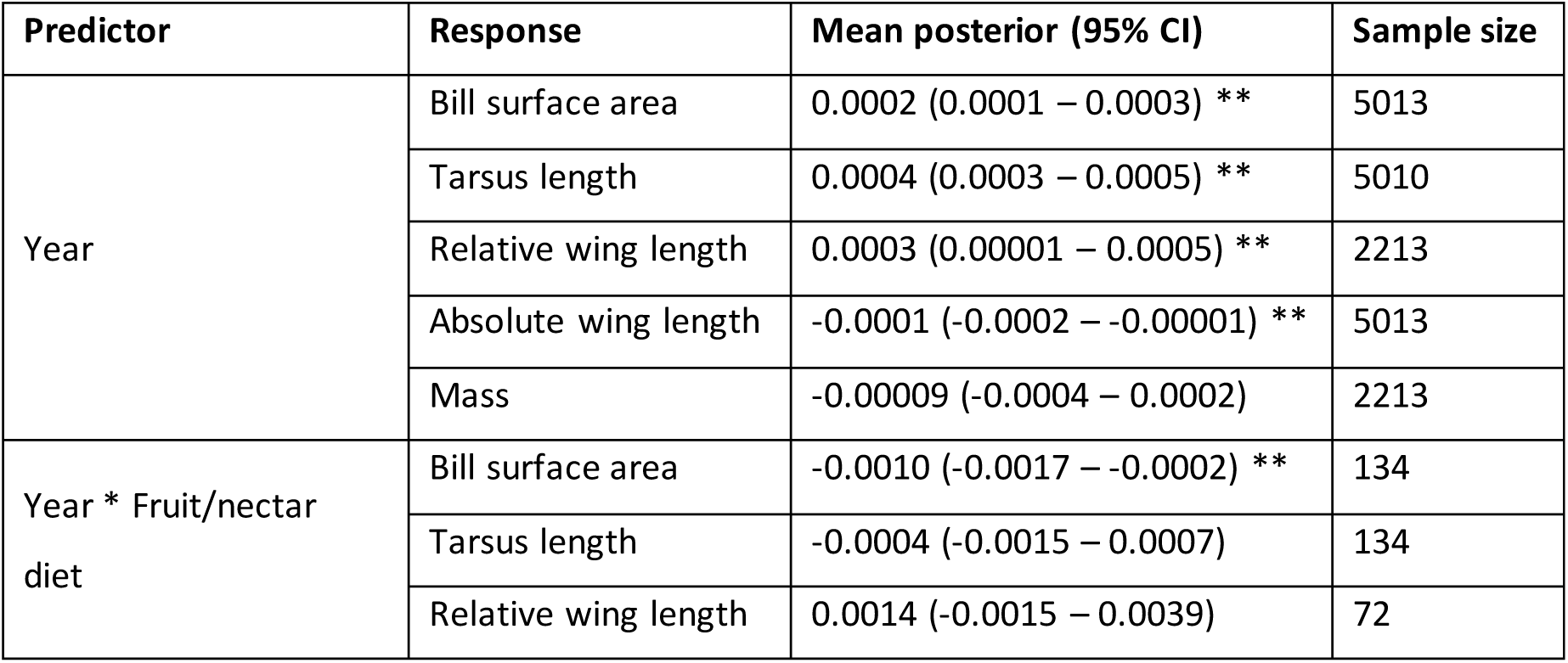
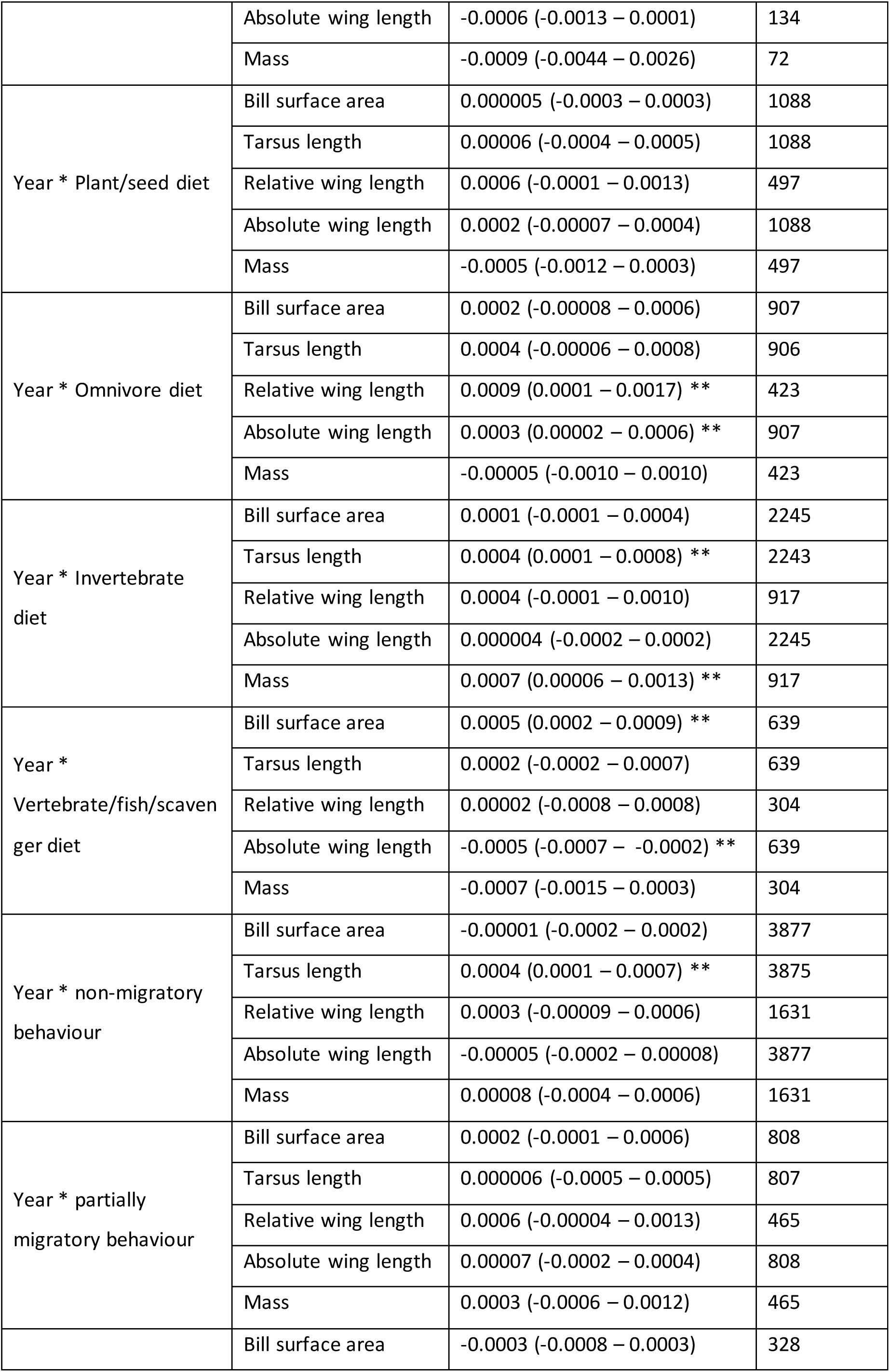
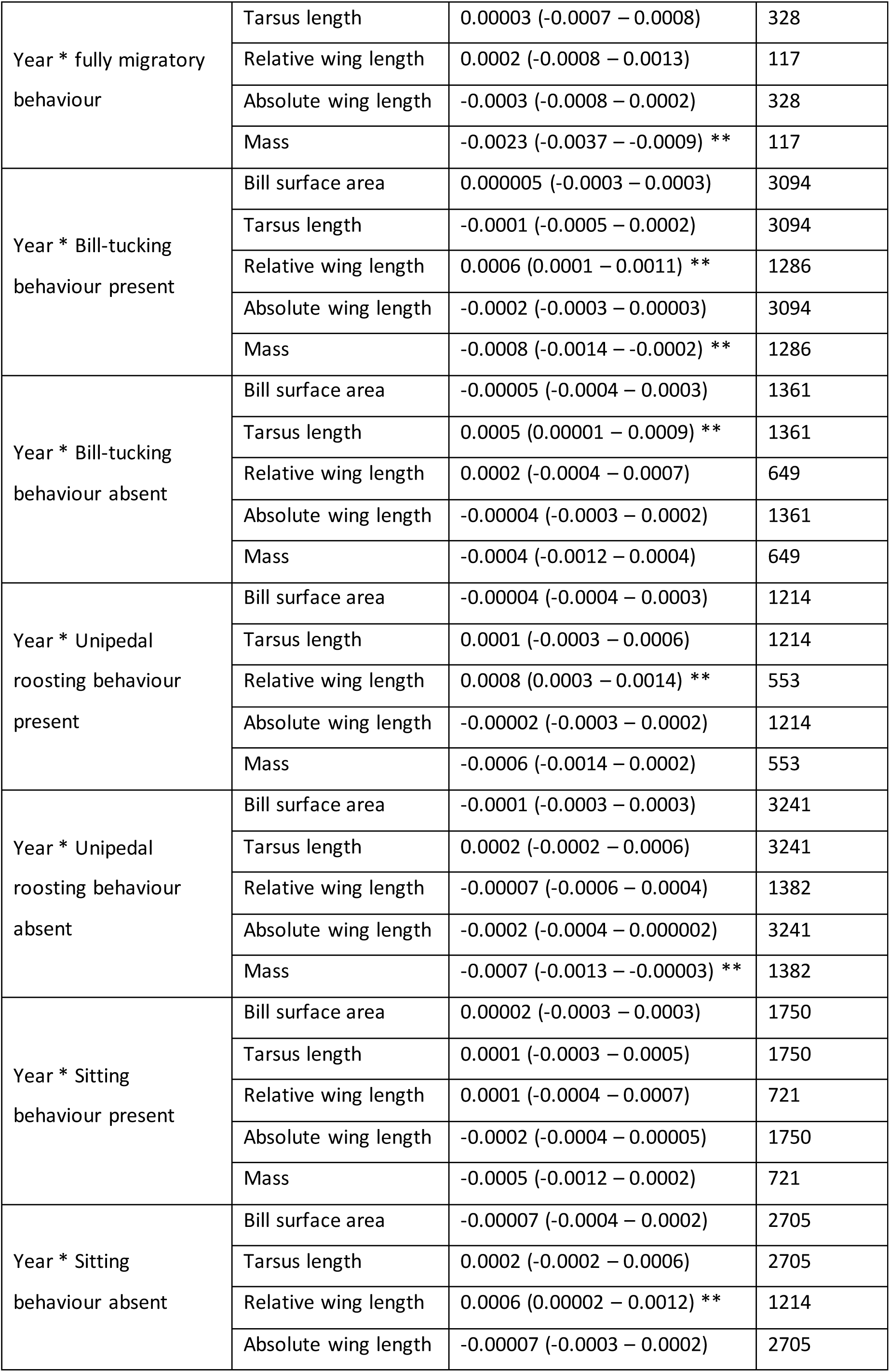
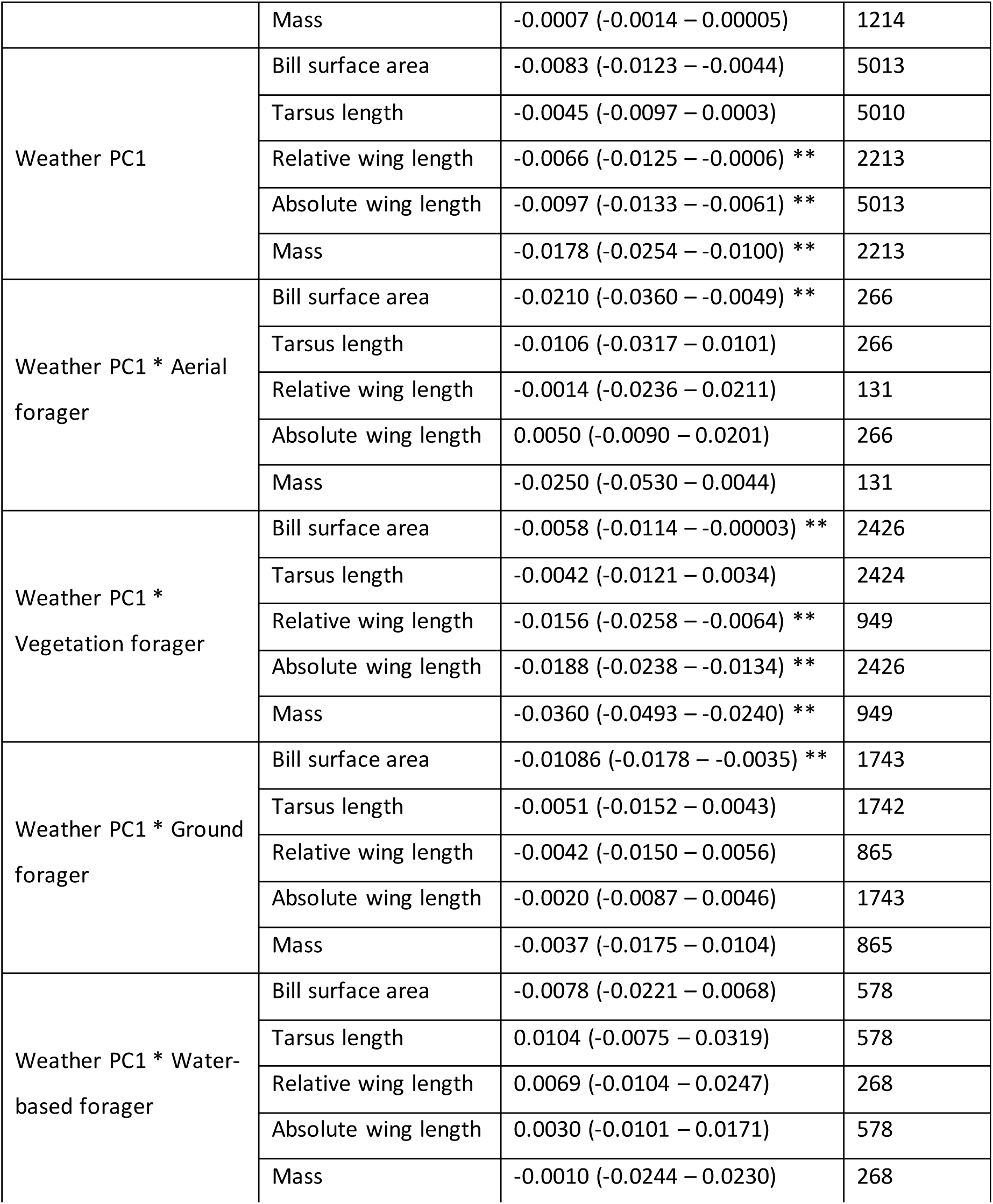
Morphological changes through time and in response to hotter weather in the 5 years preceding collection. Values reported are mean of the posterior with 95% credible intervals for each predictor and response variable, with sample sizes for that combination. Stars signify significant (i.e. 95% CI does not cross 0) effects. For categorical model predictors, values reported are marginal effect estimates.

**Table ED2.**
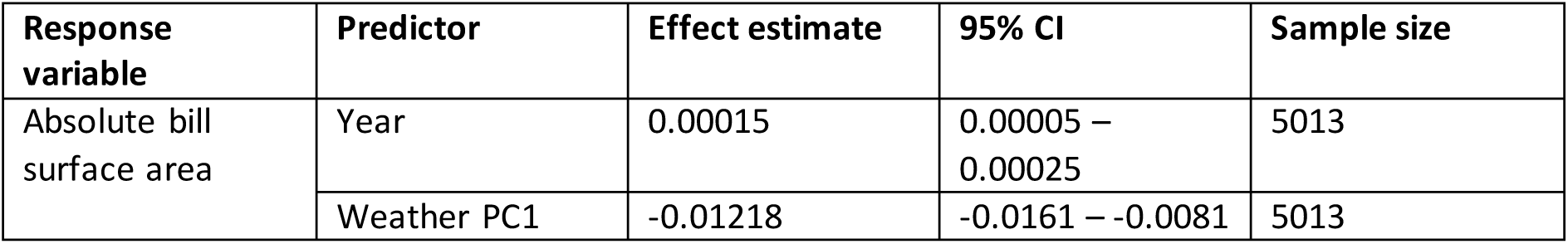
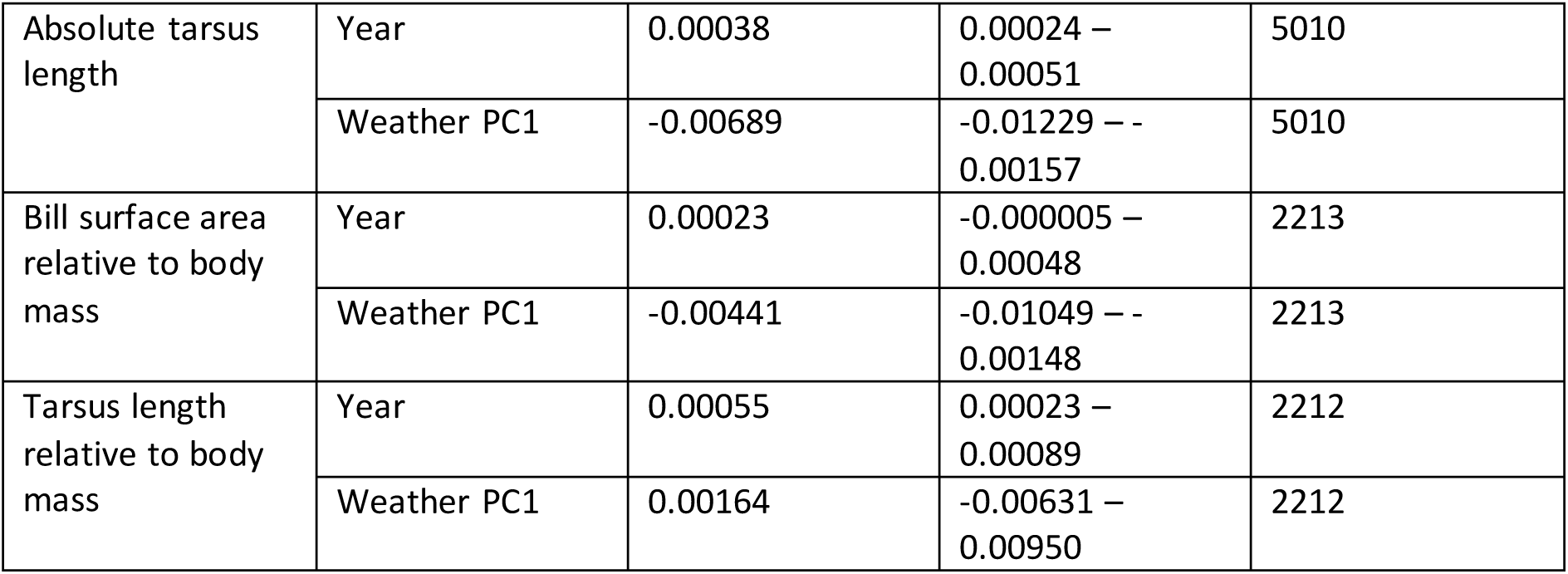
Model outputs for changes in absolute bill surface area and tarsus length, and changes in bill surface area and tarsus length relative to mass.<.

| Response variable | Predictor | Effect estimate | 95% CI | Sample size |
| --- | --- | --- | --- | --- |
| Absolute bill surface area | Year | 0.00015 | 0.00005 – 0.00025 | 5013 |
|  | Weather PC1 | -0.01218 | -0.0161 – -0.0081 | 5013 |
| Absolute tarsus length | Year | 0.00038 | 0.00024 – 0.00051 | 5010 |
|  | Weather PC1 | -0.00689 | -0.01229 – - 0.00157 | 5010 |
| Bill surface area relative to body mass | Year | 0.00023 | -0.000005 – 0.00048 | 2213 |
|  | Weather PC1 | -0.00441 | -0.01049 – - 0.00148 | 2213 |
| Tarsus length relative to body mass | Year | 0.00055 | 0.00023 – 0.00089 | 2212 |
|  | Weather PC1 | 0.00164 | -0.00631 – 0.00950 | 2212 |

## Supplementary information

**Table S1.**
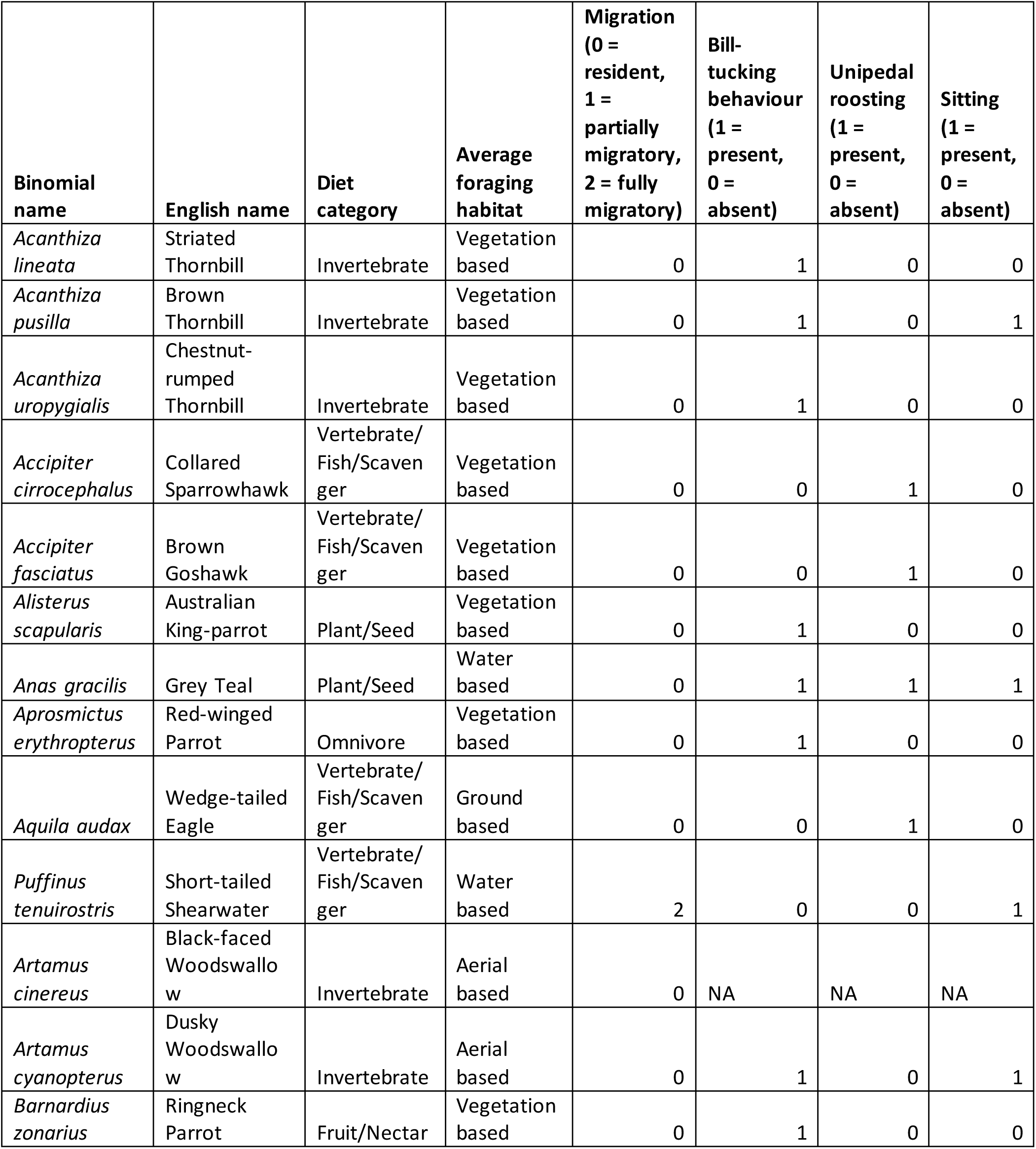
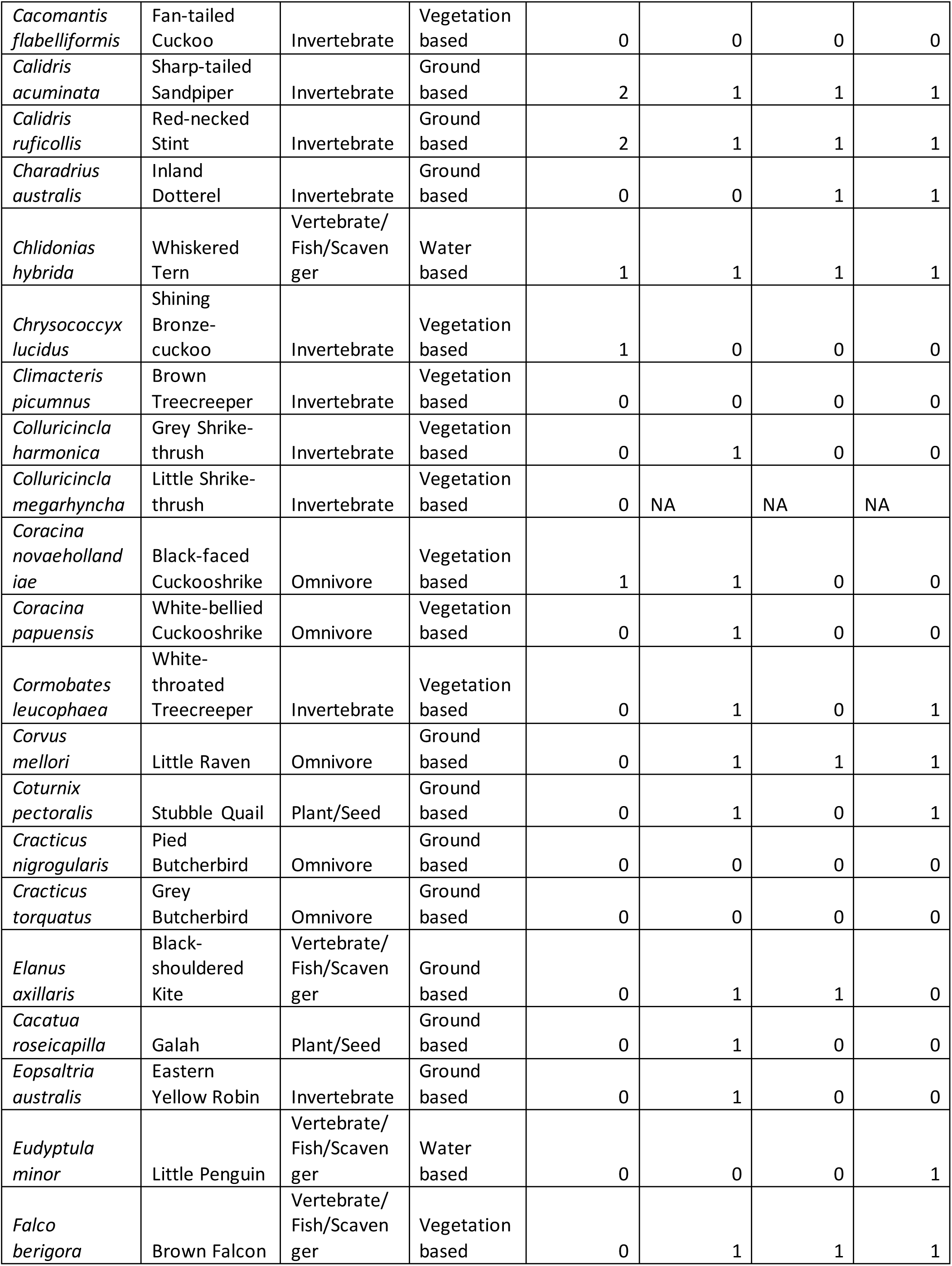
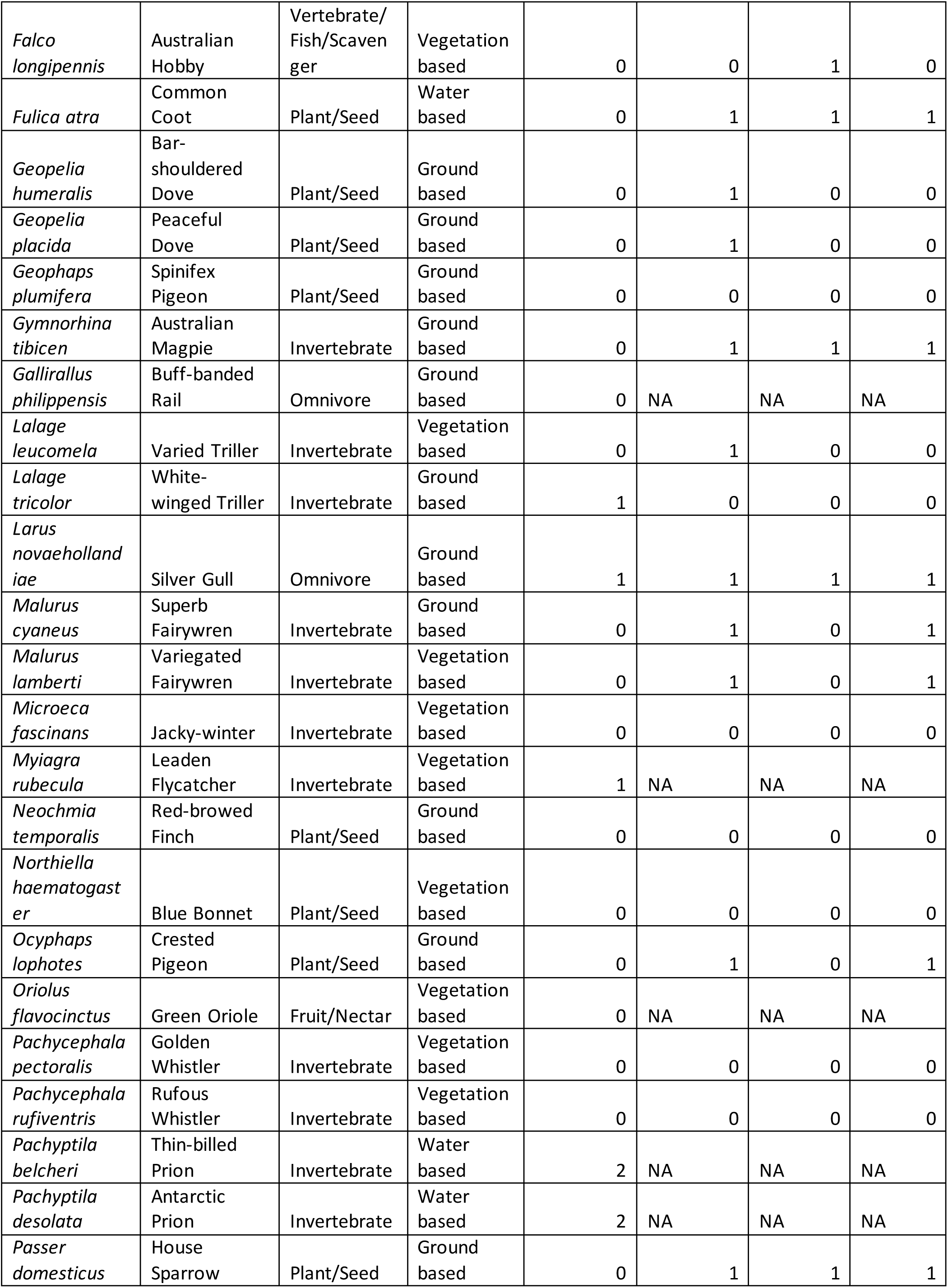
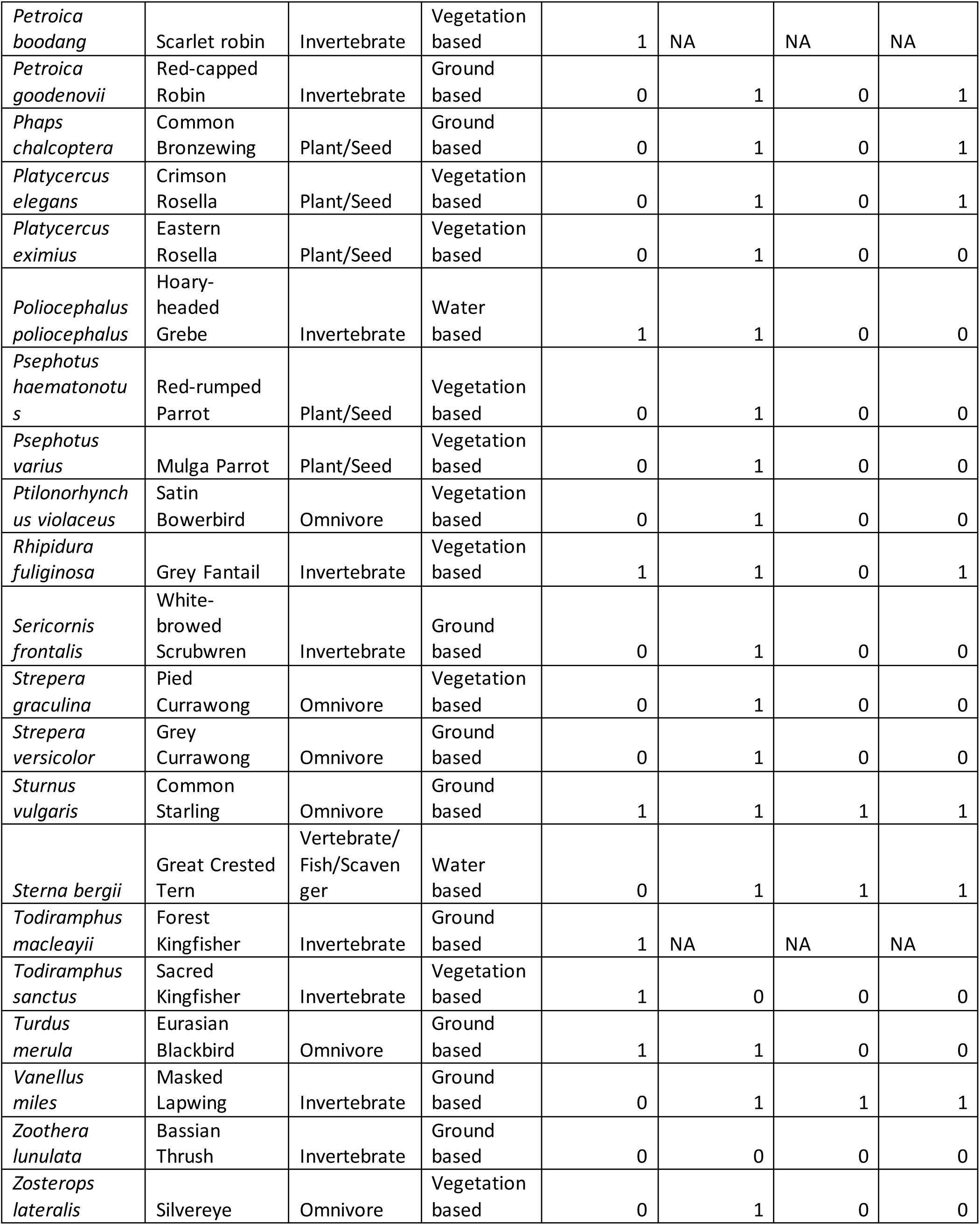
Summary of species traits collated for the analysis in this paper. Showing binomial name, English name, diet category, average foraging habitat, migratory behaviour (0=resident, 1=partial migrant, 2=full migrant), and the presence (1) or absence (0) of three thermoregulatory behaviours (bill-tucking, unipedal roosting, and sitting). NA’s represent species for whom that information was missing.

**Table S2.**
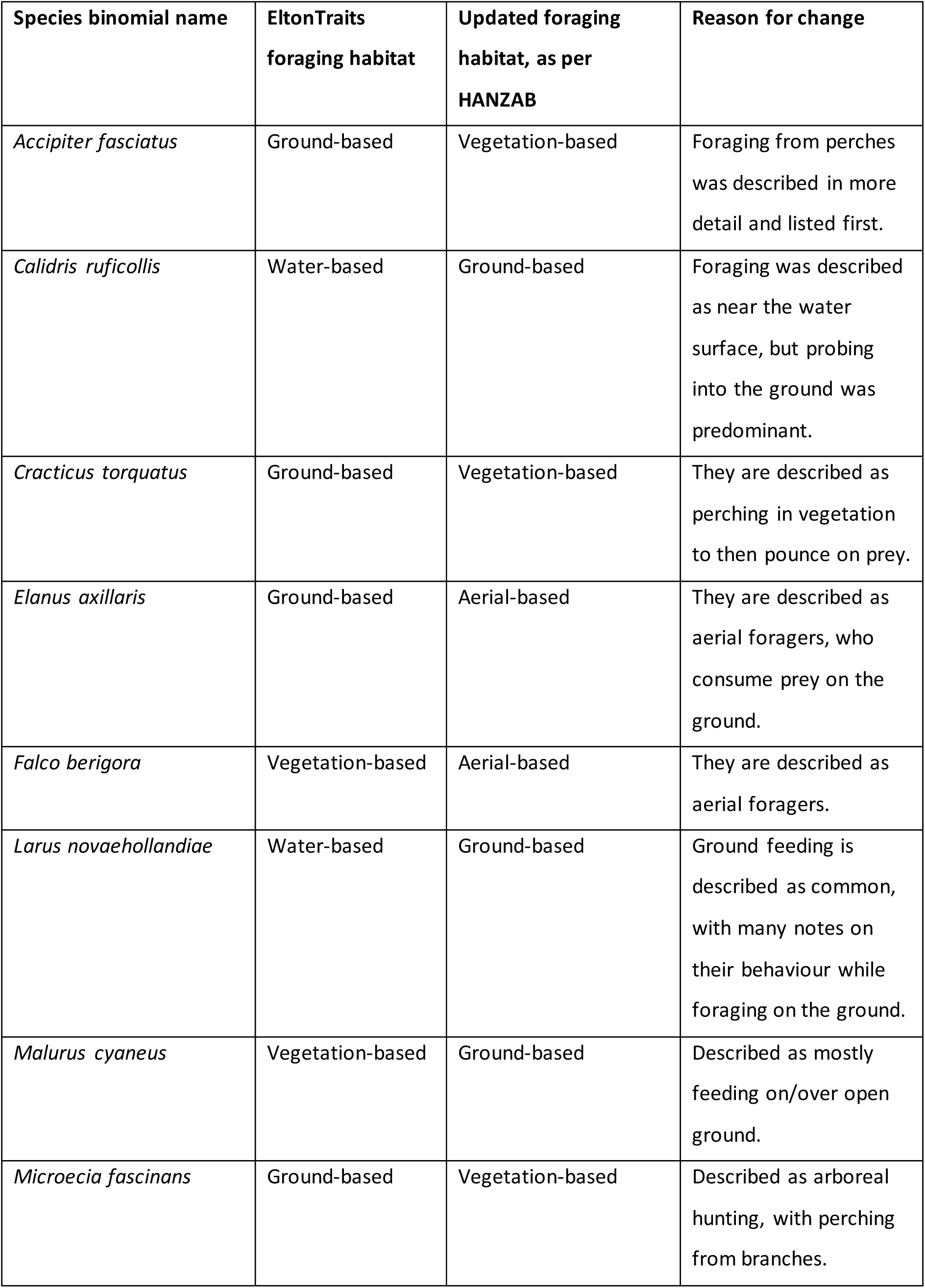
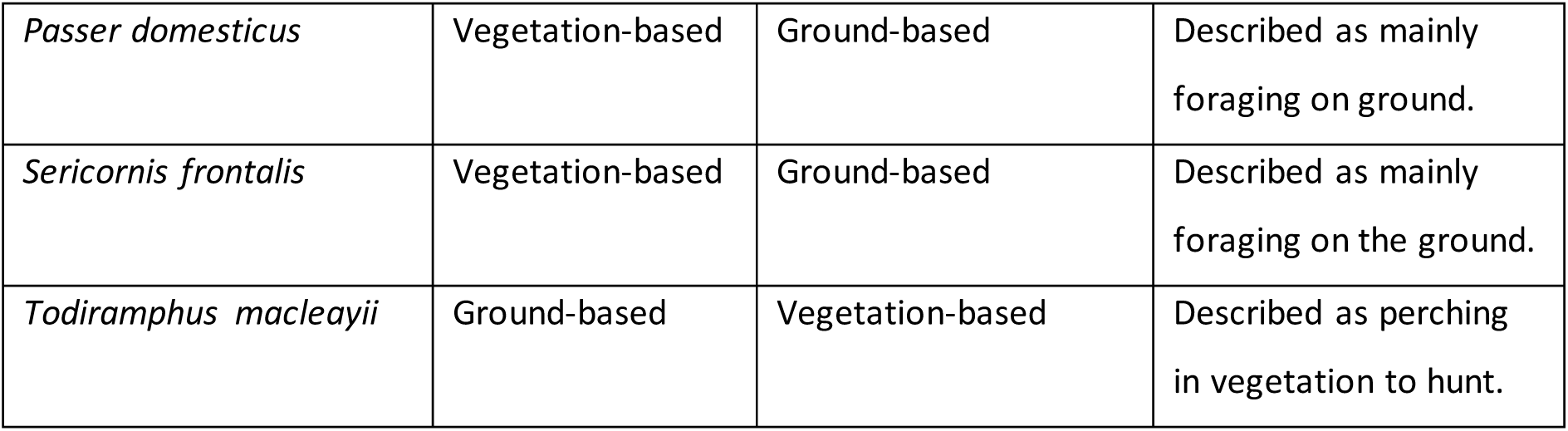
Species for whom we updated the foraging habitat taken from EltonTraits using HANZAB information. The table includes the EltonTraits designation, what it was changed to (i.e. the HANZAB entry) and the reason for the change.

**Figure S1.**
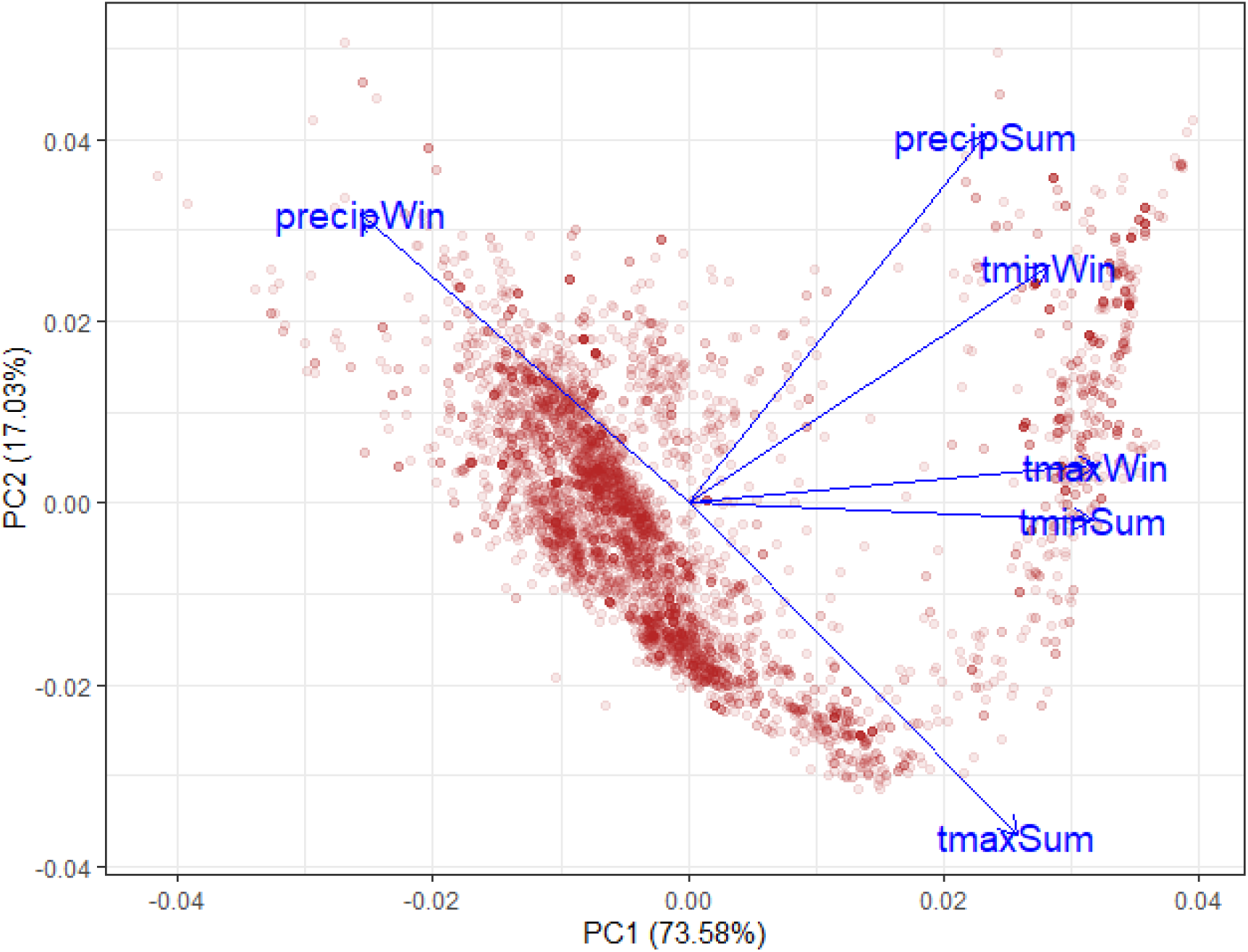
Principal component loadings from our weather PCA. PC1 was represented relatively evenly by all weather metrics. In descending order of importance, these were: maximum winter temperature (0.468), minimum summer temperature (0.462), minimum winter temperature (0.412), maximum summer temperature (0.377), winter precipitation (–0.375), and summer precipitation (0.340).

**Table S3.**
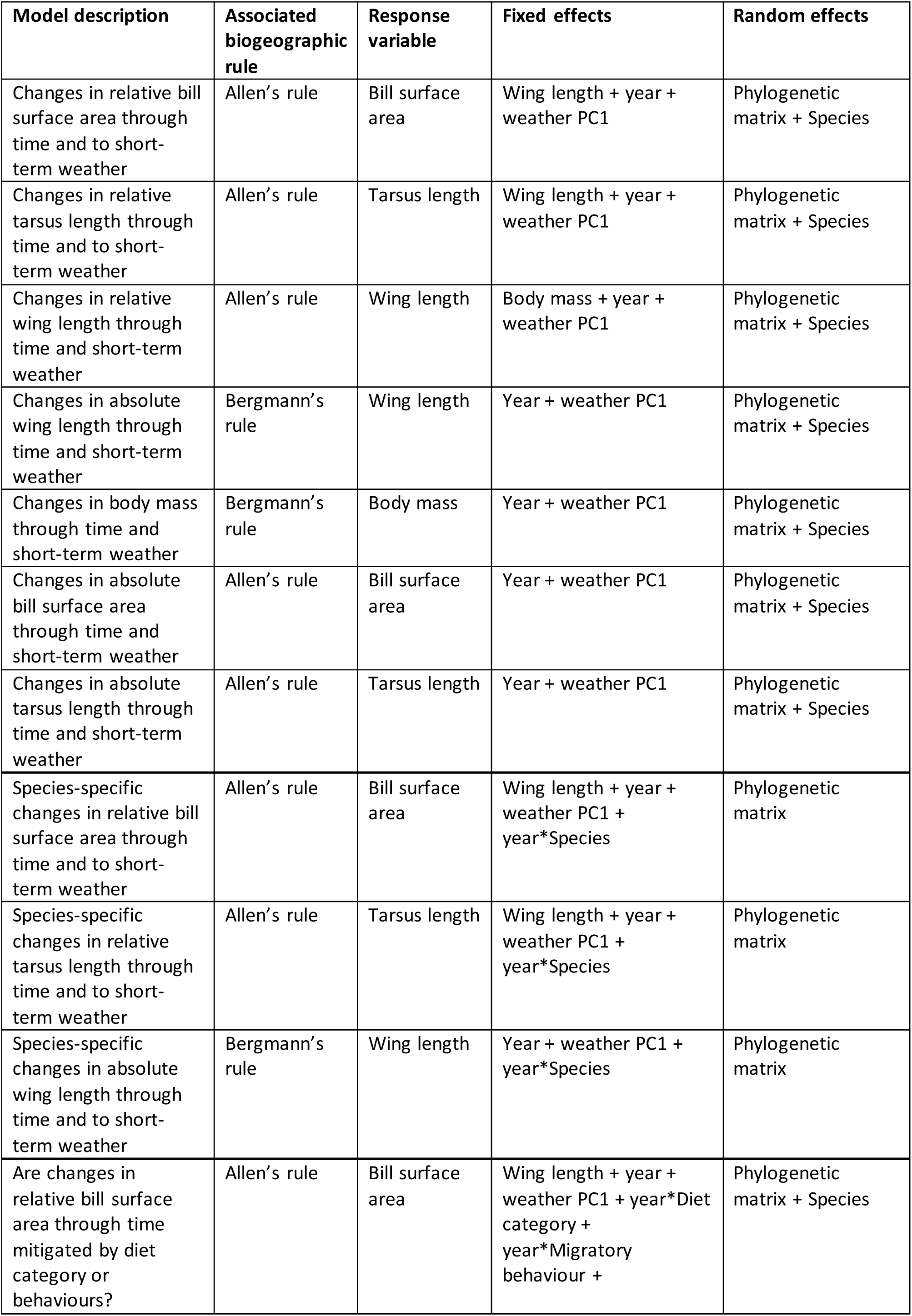
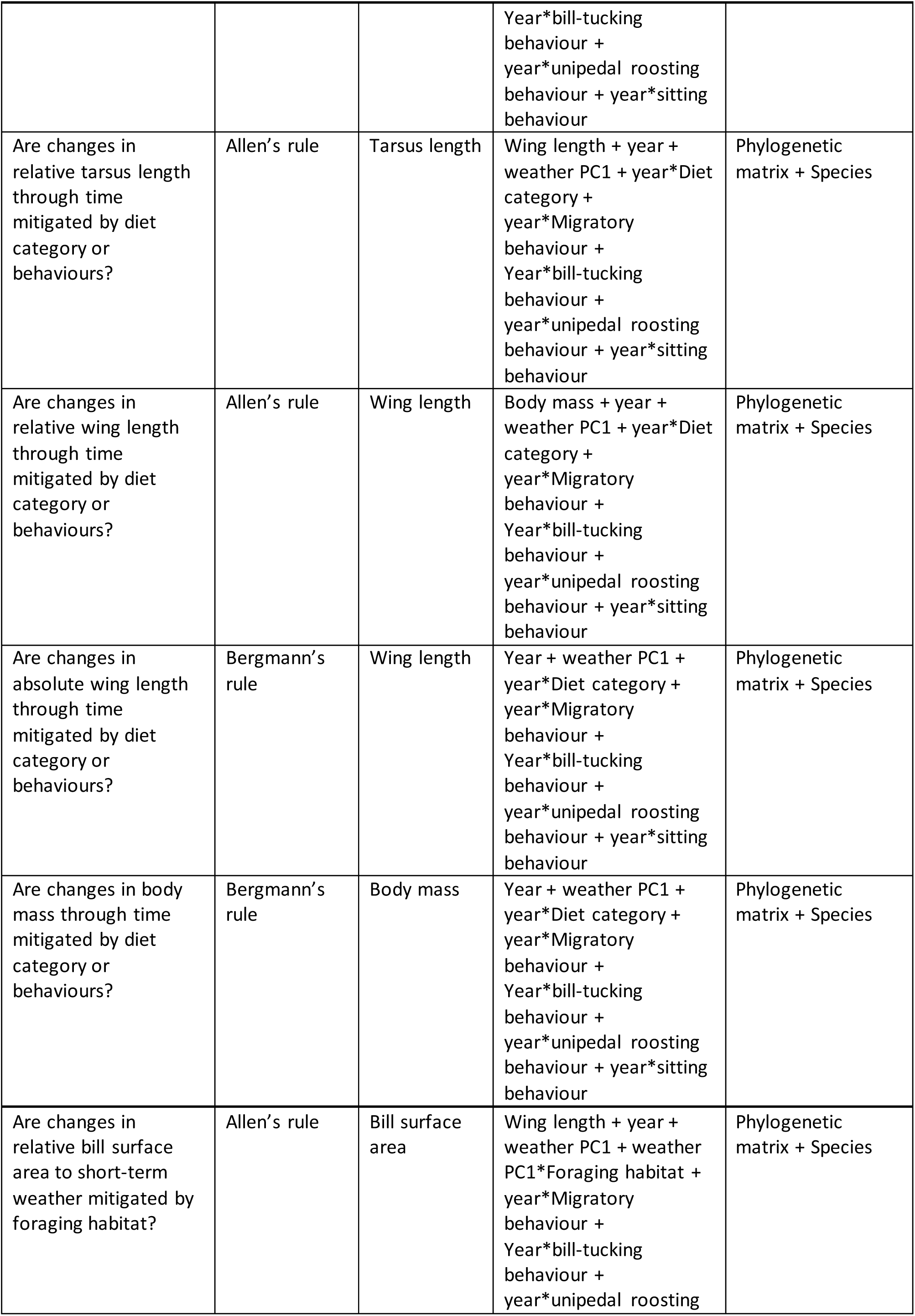
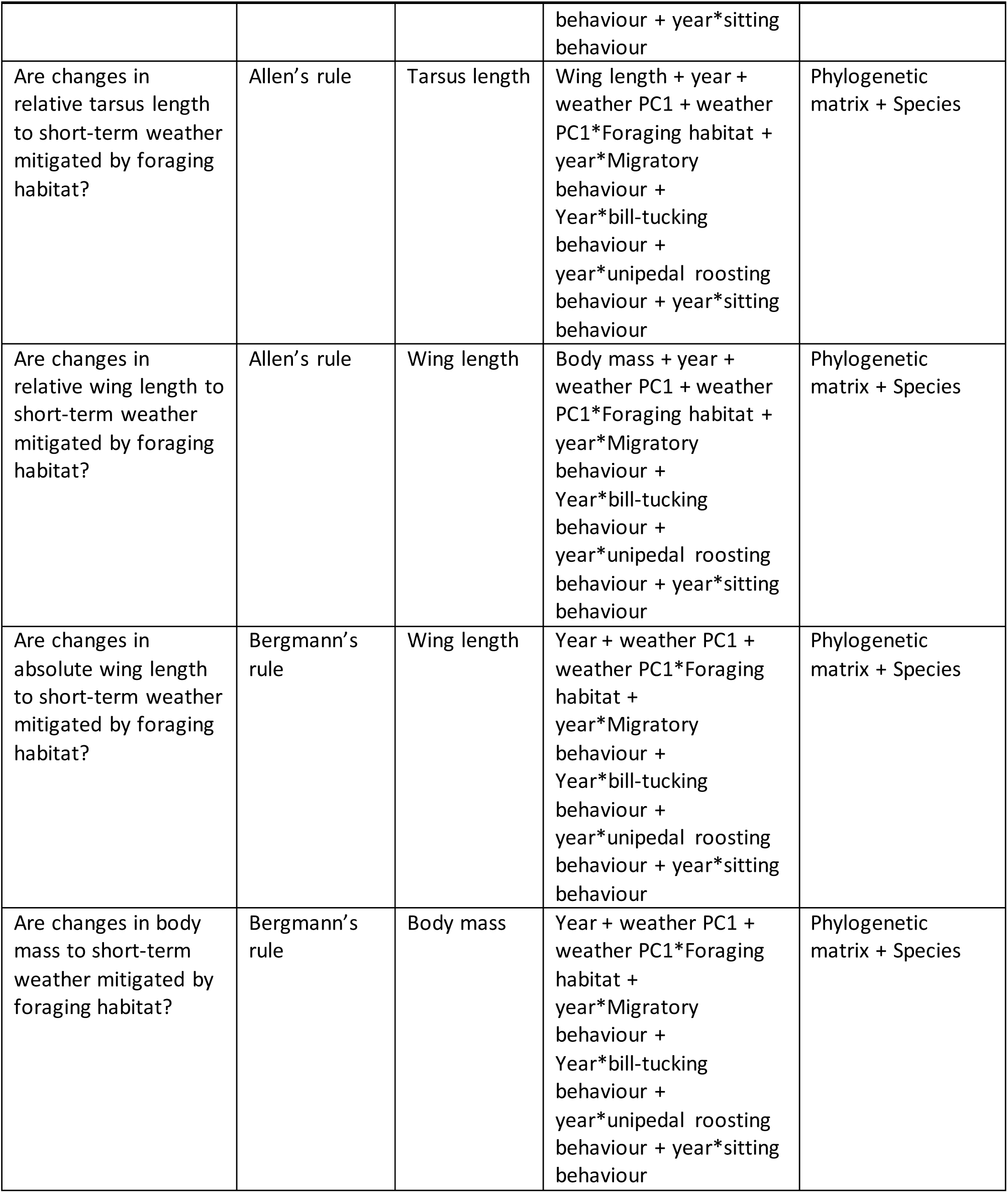
Summary of all the models run, with a description of what each is testing, the biogeographic rule it is associated with (Allen’s or Bergmann’s rule), the response variable, the fixed effects, and the random effects.

## Notes

### Competing Interest Statement

The authors have declared no competing interest.

