## Supplementary tables 1-3 and figure 1 for "Pervasive morphological responses to climate change in bird body and appendage size"

**Supplementary information**

**Table S1.** Summary of species traits collated for the analysis in this paper. Showing binomial name, English name, diet category, average foraging habitat, migratory behaviour (0=resident, 1=partial migrant, 2=full migrant), and the presence (1) or absence (0) of three thermoregulatory behaviours (bill-tucking, unipedal roosting, and sitting). NA's represent species for whom that information was missing.

| **Binomial name** | **English name** | **Diet category** | **Average foraging habitat** | **Migration (0 = resident, 1 = partially migratory, 2 = fully migratory)** | **Bill-tucking behaviour (1 = present, 0 = absent)** | **Unipedal roosting (1 = present, 0 = absent)** | **Sitting (1 = present, 0 = absent)** |
|---|---|---|---|---|---|---|---|
| *Acanthiza lineata* | Striated Thornbill | Invertebrate | Vegetation based | 0 | 1 | 0 | 0 |
| *Acanthiza pusilla* | Brown Thornbill | Invertebrate | Vegetation based | 0 | 1 | 0 | 1 |
| *Acanthiza uropygialis* | Chestnut-rumped Thornbill | Invertebrate | Vegetation based | 0 | 1 | 0 | 0 |
| *Accipiter cirrocephalus* | Collared Sparrowhawk | Vertebrate/ Fish/Scavenger | Vegetation based | 0 | 0 | 1 | 0 |
| *Accipiter fasciatus* | Brown Goshawk | Vertebrate/ Fish/Scavenger | Vegetation based | 0 | 0 | 1 | 0 |
| *Alisterus scapularis* | Australian King-parrot | Plant/Seed | Vegetation based | 0 | 1 | 0 | 0 |
| *Anas gracilis* | Grey Teal | Plant/Seed | Water based | 0 | 1 | 1 | 1 |
| *Aprosmictus erythropterus* | Red-winged Parrot | Omnivore | Vegetation based | 0 | 1 | 0 | 0 |
| *Aquila audax* | Wedge-tailed Eagle | Vertebrate/ Fish/Scavenger | Ground based | 0 | 0 | 1 | 0 |
| *Puffinus tenuirostris* | Short-tailed Shearwater | Vertebrate/ Fish/Scavenger | Water based | 2 | 0 | 0 | 1 |
| *Artamus cinereus* | Black-faced Woodswallow | Invertebrate | Aerial based | 0 | NA | NA | NA |
| *Artamus cyanopterus* | Dusky Woodswallow | Invertebrate | Aerial based | 0 | 1 | 0 | 1 |
| *Barnardius zonarius* | Ringneck Parrot | Fruit/Nectar | Vegetation based | 0 | 1 | 0 | 0 |

| | | | | | | | |
|---|---|---|---|---|---|---|---|
| *Cacomantis flabelliformis* | Fan-tailed Cuckoo | Invertebrate | Vegetation based | 0 | 0 | 0 | 0 |
| *Calidris acuminata* | Sharp-tailed Sandpiper | Invertebrate | Ground based | 2 | 1 | 1 | 1 |
| *Calidris ruficollis* | Red-necked Stint | Invertebrate | Ground based | 2 | 1 | 1 | 1 |
| *Charadrius australis* | Inland Dotterel | Invertebrate | Ground based | 0 | 0 | 1 | 1 |
| *Chlidonias hybrida* | Whiskered Tern | Vertebrate/ Fish/Scavenger | Water based | 1 | 1 | 1 | 1 |
| *Chrysococcyx lucidus* | Shining Bronze-cuckoo | Invertebrate | Vegetation based | 1 | 0 | 0 | 0 |
| *Climacteris picumnus* | Brown Treecreeper | Invertebrate | Vegetation based | 0 | 0 | 0 | 0 |
| *Colluricincla harmonica* | Grey Shrike-thrush | Invertebrate | Vegetation based | 0 | 1 | 0 | 0 |
| *Colluricincla megarhyncha* | Little Shrike-thrush | Invertebrate | Vegetation based | 0 | NA | NA | NA |
| *Coracina novaehollandiae* | Black-faced Cuckooshrike | Omnivore | Vegetation based | 1 | 1 | 0 | 0 |
| *Coracina papuensis* | White-bellied Cuckooshrike | Omnivore | Vegetation based | 0 | 1 | 0 | 0 |
| *Cormobates leucophaea* | White-throated Treecreeper | Invertebrate | Vegetation based | 0 | 1 | 0 | 1 |
| *Corvus mellori* | Little Raven | Omnivore | Ground based | 0 | 1 | 1 | 1 |
| *Coturnix pectoralis* | Stubble Quail | Plant/Seed | Ground based | 0 | 1 | 0 | 1 |
| *Cracticus nigrogularis* | Pied Butcherbird | Omnivore | Ground based | 0 | 0 | 0 | 0 |
| *Cracticus torquatus* | Grey Butcherbird | Omnivore | Ground based | 0 | 0 | 0 | 0 |
| *Elanus axillaris* | Black-shouldered Kite | Vertebrate/ Fish/Scavenger | Ground based | 0 | 1 | 1 | 0 |
| *Cacatua roseicapilla* | Galah | Plant/Seed | Ground based | 0 | 1 | 0 | 0 |
| *Eopsaltria australis* | Eastern Yellow Robin | Invertebrate | Ground based | 0 | 1 | 0 | 0 |
| *Eudyptula minor* | Little Penguin | Vertebrate/ Fish/Scavenger | Water based | 0 | 0 | 0 | 1 |
| *Falco berigora* | Brown Falcon | Vertebrate/ Fish/Scavenger | Vegetation based | 0 | 1 | 1 | 1 |

| | | | | | | | |
|---|---|---|---|---|---|---|---|
| *Falco longipennis* | Australian Hobby | Vertebrate/ Fish/Scavenger | Vegetation based | 0 | 0 | 1 | 0 |
| *Fulica atra* | Common Coot | Plant/Seed | Water based | 0 | 1 | 1 | 1 |
| *Geopelia humeralis* | Bar-shouldered Dove | Plant/Seed | Ground based | 0 | 1 | 0 | 0 |
| *Geopelia placida* | Peaceful Dove | Plant/Seed | Ground based | 0 | 1 | 0 | 0 |
| *Geophaps plumifera* | Spinifex Pigeon | Plant/Seed | Ground based | 0 | 0 | 0 | 0 |
| *Gymnorhina tibicen* | Australian Magpie | Invertebrate | Ground based | 0 | 1 | 1 | 1 |
| *Gallirallus philippensis* | Buff-banded Rail | Omnivore | Ground based | 0 | NA | NA | NA |
| *Lalage leucomela* | Varied Triller | Invertebrate | Vegetation based | 0 | 1 | 0 | 0 |
| *Lalage tricolor* | White-winged Triller | Invertebrate | Ground based | 1 | 0 | 0 | 0 |
| *Larus novaehollandiae* | Silver Gull | Omnivore | Ground based | 1 | 1 | 1 | 1 |
| *Malurus cyaneus* | Superb Fairywren | Invertebrate | Ground based | 0 | 1 | 0 | 1 |
| *Malurus lamberti* | Variegated Fairywren | Invertebrate | Vegetation based | 0 | 1 | 0 | 1 |
| *Microeca fascinans* | Jacky-winter | Invertebrate | Vegetation based | 0 | 0 | 0 | 0 |
| *Myiagra rubecula* | Leaden Flycatcher | Invertebrate | Vegetation based | 1 | NA | NA | NA |
| *Neochmia temporalis* | Red-browed Finch | Plant/Seed | Ground based | 0 | 0 | 0 | 0 |
| *Northiella haematogaster* | Blue Bonnet | Plant/Seed | Vegetation based | 0 | 0 | 0 | 0 |
| *Ocyphaps lophotes* | Crested Pigeon | Plant/Seed | Ground based | 0 | 1 | 0 | 1 |
| *Oriolus flavocinctus* | Green Oriole | Fruit/Nectar | Vegetation based | 0 | NA | NA | NA |
| *Pachycephala pectoralis* | Golden Whistler | Invertebrate | Vegetation based | 0 | 0 | 0 | 0 |
| *Pachycephala rufiventris* | Rufous Whistler | Invertebrate | Vegetation based | 0 | 0 | 0 | 0 |
| *Pachyptila belcheri* | Thin-billed Prion | Invertebrate | Water based | 2 | NA | NA | NA |
| *Pachyptila desolata* | Antarctic Prion | Invertebrate | Water based | 2 | NA | NA | NA |
| *Passer domesticus* | House Sparrow | Plant/Seed | Ground based | 0 | 1 | 1 | 1 |

| | | | | | | | |
|---|---|---|---|---|---|---|---|
| *Petroica boodang* | Scarlet robin | Invertebrate | Vegetation based | 1 | NA | NA | NA |
| *Petroica goodenovii* | Red-capped Robin | Invertebrate | Ground based | 0 | 1 | 0 | 1 |
| *Phaps chalcoptera* | Common Bronzewing | Plant/Seed | Ground based | 0 | 1 | 0 | 1 |
| *Platycercus elegans* | Crimson Rosella | Plant/Seed | Vegetation based | 0 | 1 | 0 | 1 |
| *Platycercus eximius* | Eastern Rosella | Plant/Seed | Vegetation based | 0 | 1 | 0 | 0 |
| *Poliocephalus poliocephalus* | Hoary-headed Grebe | Invertebrate | Water based | 1 | 1 | 0 | 0 |
| *Psephotus haematonotus* | Red-rumped Parrot | Plant/Seed | Vegetation based | 0 | 1 | 0 | 0 |
| *Psephotus varius* | Mulga Parrot | Plant/Seed | Vegetation based | 0 | 1 | 0 | 0 |
| *Ptilonorhynchus violaceus* | Satin Bowerbird | Omnivore | Vegetation based | 0 | 1 | 0 | 0 |
| *Rhipidura fuliginosa* | Grey Fantail | Invertebrate | Vegetation based | 1 | 1 | 0 | 1 |
| *Sericornis frontalis* | White-browed Scrubwren | Invertebrate | Ground based | 0 | 1 | 0 | 0 |
| *Strepera graculina* | Pied Currawong | Omnivore | Vegetation based | 0 | 1 | 0 | 0 |
| *Strepera versicolor* | Grey Currawong | Omnivore | Ground based | 0 | 1 | 0 | 0 |
| *Sturnus vulgaris* | Common Starling | Omnivore | Ground based | 1 | 1 | 1 | 1 |
| *Sterna bergii* | Great Crested Tern | Vertebrate/ Fish/Scavenger | Water based | 0 | 1 | 1 | 1 |
| *Todiramphus macleayii* | Forest Kingfisher | Invertebrate | Ground based | 1 | NA | NA | NA |
| *Todiramphus sanctus* | Sacred Kingfisher | Invertebrate | Vegetation based | 1 | 0 | 0 | 0 |
| *Turdus merula* | Eurasian Blackbird | Omnivore | Ground based | 1 | 1 | 0 | 0 |
| *Vanellus miles* | Masked Lapwing | Invertebrate | Ground based | 0 | 1 | 1 | 1 |
| *Zoothera lunulata* | Bassian Thrush | Invertebrate | Ground based | 0 | 0 | 0 | 0 |
| *Zosterops lateralis* | Silvereye | Omnivore | Vegetation based | 0 | 1 | 0 | 0 |

**Table S2.** Species for whom we updated the foraging habitat taken from EltonTraits using HANZAB information. The table includes the EltonTraits designation, what it was changed to (i.e. the HANZAB entry) and the reason for the change.

| **Species binomial name** | **EltonTraits foraging habitat** | **Updated foraging habitat, as per HANZAB** | **Reason for change** |
|---|---|---|---|
| *Accipiter fasciatus* | Ground-based | Vegetation-based | Foraging from perches was described in more detail and listed first. |
| *Calidris ruficollis* | Water-based | Ground-based | Foraging was described as near the water surface, but probing into the ground was predominant. |
| *Cracticus torquatus* | Ground-based | Vegetation-based | They are described as perching in vegetation to then pounce on prey. |
| *Elanus axillaris* | Ground-based | Aerial-based | They are described as aerial foragers, who consume prey on the ground. |
| *Falco berigora* | Vegetation-based | Aerial-based | They are described as aerial foragers. |
| *Larus novaehollandiae* | Water-based | Ground-based | Ground feeding is described as common, with many notes on their behaviour while foraging on the ground. |
| *Malurus cyaneus* | Vegetation-based | Ground-based | Described as mostly feeding on/over open ground. |
| *Microecia fascinans* | Ground-based | Vegetation-based | Described as arboreal hunting, with perching from branches. |

| | | | |
|---|---|---|---|
| *Passer domesticus* | Vegetation-based | Ground-based | Described as mainly foraging on ground. |
| *Sericornis frontalis* | Vegetation-based | Ground-based | Described as mainly foraging on the ground. |
| *Todiramphus macleayii* | Ground-based | Vegetation-based | Described as perching in vegetation to hunt. |

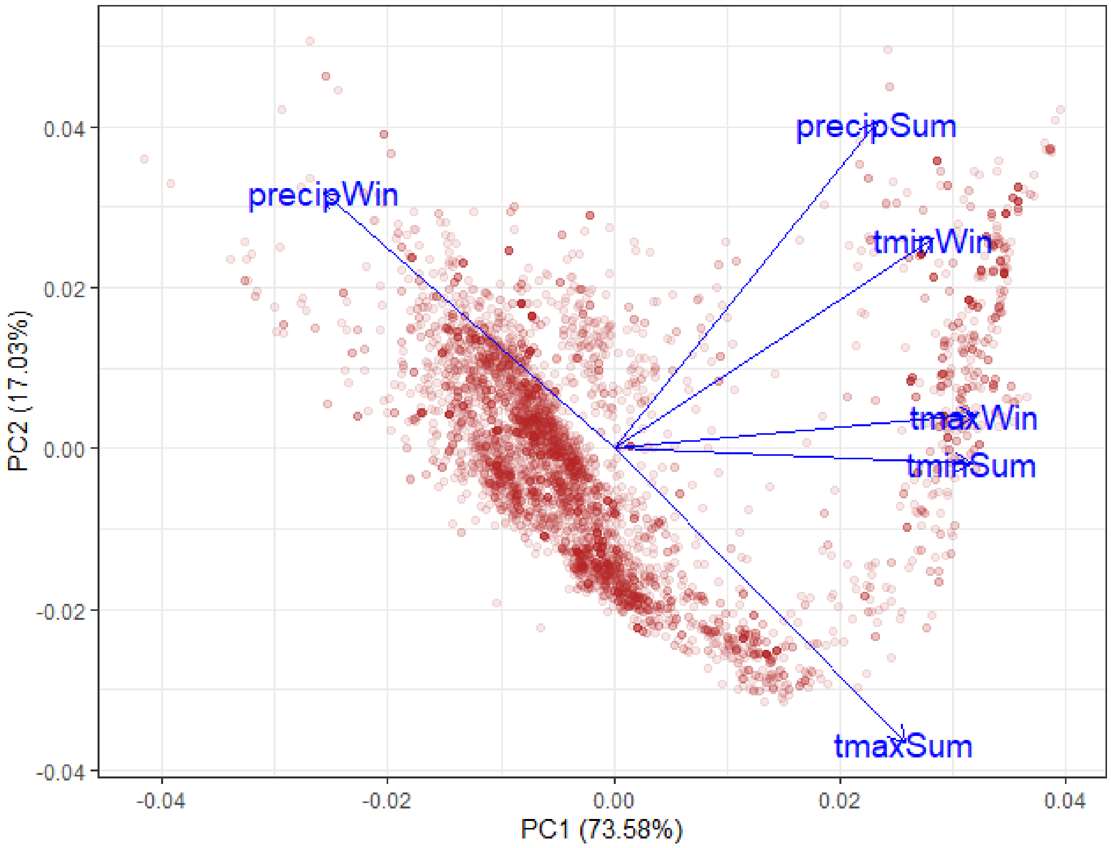


**Figure S1.** Principal component loadings from our weather PCA. PC1 was represented relatively evenly by all weather metrics. In descending order of importance, these were: maximum winter temperature (0.468), minimum summer temperature (0.462), minimum winter temperature (0.412), maximum summer temperature (0.377), winter precipitation (-0.375), and summer precipitation (0.340).

**Table S3.** Summary of all the models run, with a description of what each is testing, the biogeographic rule it is associated with (Allen's or Bergmann's rule), the response variable, the fixed effects, and the random effects

| Model description | Associated biogeographic rule | Response variable | Fixed effects | Random effects |
|---|---|---|---|---|
| Changes in relative bill surface area through time and to short-term weather | Allen's rule | Bill surface area | Wing length + year + weather PC1 | Phylogenetic matrix + Species |
| Changes in relative tarsus length through time and to short-term weather | Allen's rule | Tarsus length | Wing length + year + weather PC1 | Phylogenetic matrix + Species |
| Changes in relative wing length through time and short-term weather | Allen's rule | Wing length | Body mass + year + weather PC1 | Phylogenetic matrix + Species |
| Changes in absolute wing length through time and short-term weather | Bergmann's rule | Wing length | Year + weather PC1 | Phylogenetic matrix + Species |
| Changes in body mass through time and short-term weather | Bergmann's rule | Body mass | Year + weather PC1 | Phylogenetic matrix + Species |
| Changes in absolute bill surface area through time and short-term weather | Allen's rule | Bill surface area | Year + weather PC1 | Phylogenetic matrix + Species |
| Changes in absolute tarsus length through time and short-term weather | Allen's rule | Tarsus length | Year + weather PC1 | Phylogenetic matrix + Species |
| Species-specific changes in relative bill surface area through time and to short-term weather | Allen's rule | Bill surface area | Wing length + year + weather PC1 + year*Species | Phylogenetic matrix |
| Species-specific changes in relative tarsus length through time and to short-term weather | Allen's rule | Tarsus length | Wing length + year + weather PC1 + year*Species | Phylogenetic matrix |
| Species-specific changes in absolute wing length through time and to short-term weather | Bergmann's rule | Wing length | Year + weather PC1 + year*Species | Phylogenetic matrix |
| Are changes in relative bill surface area through time mitigated by diet category or behaviours? | Allen's rule | Bill surface area | Wing length + year + weather PC1 + year*Diet category + year*Migratory behaviour + | Phylogenetic matrix + Species |

| | | | | |
|---|---|---|---|---|
| | | | Year*bill-tucking behaviour + year*unipedal roosting behaviour + year*sitting behaviour | |
| Are changes in relative tarsus length through time mitigated by diet category or behaviours? | Allen’s rule | Tarsus length | Wing length + year + weather PC1 + year*Diet category + year*Migratory behaviour + Year*bill-tucking behaviour + year*unipedal roosting behaviour + year*sitting behaviour | Phylogenetic matrix + Species |
| Are changes in relative wing length through time mitigated by diet category or behaviours? | Allen’s rule | Wing length | Body mass + year + weather PC1 + year*Diet category + year*Migratory behaviour + Year*bill-tucking behaviour + year*unipedal roosting behaviour + year*sitting behaviour | Phylogenetic matrix + Species |
| Are changes in absolute wing length through time mitigated by diet category or behaviours? | Bergmann’s rule | Wing length | Year + weather PC1 + year*Diet category + year*Migratory behaviour + Year*bill-tucking behaviour + year*unipedal roosting behaviour + year*sitting behaviour | Phylogenetic matrix + Species |
| Are changes in body mass through time mitigated by diet category or behaviours? | Bergmann’s rule | Body mass | Year + weather PC1 + year*Diet category + year*Migratory behaviour + Year*bill-tucking behaviour + year*unipedal roosting behaviour + year*sitting behaviour | Phylogenetic matrix + Species |
| Are changes in relative bill surface area to short-term weather mitigated by foraging habitat? | Allen’s rule | Bill surface area | Wing length + year + weather PC1 + weather PC1*Foraging habitat + year*Migratory behaviour + Year*bill-tucking behaviour + year*unipedal roosting | Phylogenetic matrix + Species |

| | | | | |
|---|---|---|---|---|
| | | | behaviour + year*sitting behaviour | |
| Are changes in relative tarsus length to short-term weather mitigated by foraging habitat? | Allen's rule | Tarsus length | Wing length + year + weather PC1 + weather PC1*Foraging habitat + year*Migratory behaviour + Year*bill-tucking behaviour + year*unipedal roosting behaviour + year*sitting behaviour | Phylogenetic matrix + Species |
| Are changes in relative wing length to short-term weather mitigated by foraging habitat? | Allen's rule | Wing length | Body mass + year + weather PC1 + weather PC1*Foraging habitat + year*Migratory behaviour + Year*bill-tucking behaviour + year*unipedal roosting behaviour + year*sitting behaviour | Phylogenetic matrix + Species |
| Are changes in absolute wing length to short-term weather mitigated by foraging habitat? | Bergmann's rule | Wing length | Year + weather PC1 + weather PC1*Foraging habitat + year*Migratory behaviour + Year*bill-tucking behaviour + year*unipedal roosting behaviour + year*sitting behaviour | Phylogenetic matrix + Species |
| Are changes in body mass to short-term weather mitigated by foraging habitat? | Bergmann's rule | Body mass | Year + weather PC1 + weather PC1*Foraging habitat + year*Migratory behaviour + Year*bill-tucking behaviour + year*unipedal roosting behaviour + year*sitting behaviour | Phylogenetic matrix + Species |
